# Ureides are similarly accumulated in response to UV-C irradiation and wound but differently remobilized during recovery in *Arabidopsis* leaves

**DOI:** 10.1101/2021.06.29.450374

**Authors:** Aigerim Soltabayeva, Aizat Bekturova, Assylay Kurmanbayeva, Dinara Oshanova, Zhadyrassyn Nurbekova, Sudhakar Srivastava, Dominic Standing, Moshe Sagi

**Affiliations:** Biology Department, School of Science and Humanities, Nazarbayev University, Nur Sultan, Z05H0P9, Kazakhstan; The Albert Katz International School for Desert Studies, The Jacob Blaustein Institutes for Desert Research, Ben-Gurion University of the Negev, Sede Boqer Campus, 8499000, Israel; Jacob Blaustein Center for Scientific Cooperation, The Jacob Blaustein Institutes for Desert Research, Ben-Gurion University of the Negev, Sede Boqer Campus, 8499000, Israel; The Albert Katz Department of Dryland Biotechnologies, French Associates Institute for Agriculture and Biotechnology of Dryland, The Jacob Blaustein Institutes for Desert Research, Ben-Gurion University of the Negev, Sede Boqer Campus, 8499000, Israel

**Author notes:** Author for correspondence **Address for correspondence** Moshe Sagi, The Albert Katz Department of Dryland Biotechnologies, French Associates Institute for Agriculture and Biotechnology of Dryland, The Jacob Blaustein Institutes for Desert Research, Ben-Gurion University of the Negev, Sede Boqer Campus, 8499000, Israel. **Email addresses of contributing authors** Aigerim Soltabayeva, Aizat Bekturova, Assylay Kurmanbayeva, Dinara Oshanova, Zhadyrassyn Nurbekova, Sudhakar Srivastava, Dominic Standing.

**Keywords:** *Arabidopsis*, purine catabolism, senescence, ureide, UV-C, wound, xanthine dehydrogenase

## Abstract

To examine a role of purine degraded metabolites in response to wounding or UV-C stress, the *Arabidopsis* wild-type and *Atxdh1* KO mutants, defective in xanthine dehydrogenase1 (XDH1), were exposed to wounding and UV-C irradiation stress. In *Atxdh1* mutant, wounding or UV-C stresses resulted in lower fresh-weight, increased senescence symptoms and higher tissue cell death rate compared to Wild-type. Additionally, Wild-type exhibited lower levels of oxidative stress indicators; reactive oxygen species and malondialdehyde than *Atxdh1* mutant leaves. Notably, purine degradation transcripts and proteins were orchestrated to lead to enhanced ureide levels in Wild-type leaves 24 h after applying UV-C or wound stress. Yet, different remobilization of the accumulated ureides was noticed 72 h after stresses application. In plants treated with UV-C the allantoin level was highest in young leaves, whereas in wounded plants it was lowest in the young leaves, accumulated mainly in the middle and wounded leaves. The results indicate that in UV-C treated Wild-type, during the recovery period from stress, ureides are remobilized from the lower older leaves to support young leaf growth. In contrast, after wounding, the ureides are remobilized to the young leaves, yet more to the middle wounded leaves, to function as antioxidants and/or healing agents.

**Highlight:** UV-C and wound triggers purine degradation in old and damaged leaves to increase ureides accumulation in stress dependent rate. Impairment in purine degradation results in premature senescence in leaves.

## Introduction

Allantoin, a nitrogen-rich compound, occurs in plants as an intermediary metabolite of purine catabolism. It was shown in legumes that symbiotically fixed nitrogen is degraded to the ureides; allantoin and allantoate, and xylem transported to leaves (Smith and Atkins, 2002). In non-legume plants, purine was also shown to function as a N source, recycled and remobilized after the oxidation of xanthine and ureide hydrolysis releasing four molecule of ammonia (Werner and Witte, 2011). Ureides were shown to prevent premature senescence in old leaves of plants grown under N starvation (Soltabayeva *et al.*, 2018) as well as acting as antioxidants, alleviating the tissue damage and cell death caused by extended dark induced reactive oxygen species [ROS (Brychkova *et al.*, 2008*a*)]. UV-C and wounding stresses were shown to result in superoxide enhancement and damage tissue (Doke *et al.*, 1994), as well as hydrogen peroxide, generated locally and systemically (Orozco-Cardenas and Ryan, 1999). The possible role of purine degradation products in plant response to stresses such as drought, high salinity, and pathogen infection, which cause the production of ROS, was suggested to correlate with enhancement of endogenous allantoin levels (Montalbini, 1991; Sagi *et al.*, 1998; Alamillo *et al.*, 2010; Kanani *et al.*, 2010). Drought stress and abscisic acid enhancement was shown to induce transcript levels of XDH1 and ROS production in tomato leaves (Yesbergenova *et al.*, 2005). Plant XDH has a significant contribution to the plant redox milieu (30 to 60%) under normal and stressed conditions (Yesbergenova *et al.*, 2005; Brychkova *et al.*, 2008*a*), enhancing superoxide (O_2_^−^) generation by transferring electrons to oxygen, either from xanthine via the molybdenum cofactor catalytic center, or from NADH via the FAD domain (Yesbergenova *et al.*, 2005; Zarepour *et al.*, 2010; Werner and Witte, 2011) and references therein).

Analysis of online microarray data (GENEVESTIGATOR; https://www.genevestigator.ethz.ch) of the ureide generation genes urate oxidase (*UO*) and *XDH1* and the degradation genes allantoinase *(ALN)* and allantoicase (*AAH*) in leaves of *Arabidopsis* plants exposed to various stresses exhibited two major patterns of purine catabolism orchestration that result in the degradation or the accumulation of ureides. Stimuli such as nitrate starvation, P deficiency, drought, hypoxia, and oxidative stress, upregulated *ALN* and/or *AAH* expression and thus may result in the degradation of ureides. In contrast, stimuli such as high light, UV-B, biotic stress (*Pseudomonas syringae*), salicylic acid treatment, night extension, heat, osmotic and salt stress that down regulated *ALN* and/or *AAH* expression (Supplementary Fig. S1), may result in the accumulation of ureides. Accordingly, extended dark stress, cadmium toxicity, salt stress and high irradiance indeed led to the accumulation of ureides as the result of enhancing ureide biosynthesis gene (*XDH1* and/or *UOX*) product and repressing the ureide degradation genes (*ALN* and/or *AAH*) product (Brychkova *et al.*, 2008*a*; Irani and Todd, 2016; Nourimand and Todd, 2016; Irani *et al.*, 2018). Additionally, the opposite type of ureides regulation, i.e the reduction of allantoin content by activating the *ALN* and *AAH* gene was indeed shown under nitrate starvation (Soltabayeva *et al.*, 2018). These data indicate that the purine network genes are controlled either to degrade allantoin likely to provide nitrogen or accumulate to protect as antioxidant against oxidative stress (Brychkova *et al.*, 2008*a*,*b*; Soltabayeva *et al.*, 2018) (Supplementary Fig. S2). Yet, it remains unclear how the purine degradation pathway contributes to the stress protection mechanisms during the recovery period from the stress applied.

Exposing wild-type and knockout mutants *Atxdh1,* impaired in xanthine dehydrogenase (*XDH1*) expression, to local wounding and UV-C irradiation stress revealed that purine degradation genes and their products in both UV-C and wound leaves were orchestrated on transcript and protein levels, resulting in enhanced ureides in WT leaves 24 h after applying the stress. Notably, different remobilization of the accumulated ureides was noticed 72 h after the stress application. In plants treated with UV-C the allantoin level was highest in young leaves, whereas in wounded plants it was lowest in the young leaves, accumulating mainly in the middle and wounded leaves.

## Materials and methods

### Plant Material, Growth Conditions and treatment applied

*Arabidopsis thaliana* (L.) Heynh wild-type and mutants used in the current study were of the Col-0 background. The following homozygous T-DNA-inserted mutants were employed: *Atxdh1* (GABI_049D04, SALK_148366; accession no. At4g34890) described previously by us (Yesbergenova *et al*., 2005; Brychkova *et al*., 2008); *Ataln* (SALK_ 013427; accession no. At4g04955) as shown before (Todd and Polacco, 2006; Watanabe *et al.*, 2014)); and *Ataah* (SALK_112631; accession no. At4g20070) as described previously (Todd and Polacco, 2006).

Seeds were surface-sterilized in 80% alcohol for 2 min, washed three times in sterile water and sown on one-half strength Murashige and Skoog (0.5 MS) agar plates (Murashige and Skoog, 1962). The plates were placed at 4 °C for 3 days to synchronize germination, and then were transferred to a controlled growth room at 22°C, 14/10 h light/dark photoperiod and light intensity of 150 μE m^−2^ s ^−1^. Six-day-old seedlings were transferred to a 1:1 mixture of perlite and soil. 25 days post germination plants were exposed to mechanical wound or UV-C of middle leaves (fully exposed leaves, 5-10^th^ from bottom). Mechanical wounding was applied with forceps in the middle of leaves perpendicular to main vein. The wound was applied once per leaf in all the experiments, but in an Immunoblot analysis we additionally tested the double wounding effect in comparison to one wounding. For the UV-C experiment, plants were irradiated with 0.150 Joule/m^2^ UV-C (λ=254 nm) using a CL 508G UV cross-linker and placed immediately in the growth room for recovery for 3 days. The H_2_O_2_ production was visualized by staining the leaf discs with 3,3′-diaminobenzidine (DAB) and superoxide generation was assessed by nitroblue tetrazolium (NBT) staining 6h after wounding and 24 hours after the exposure to UV-C stress. Leaf discs were photographed after boiling in 80% ethanol for 10 min. Tissue death was visualized in leaves after 24 hours of wounding, by lactophenol–trypan blue staining, followed by de-staining in saturated chloral hydrate (Koch and Slusarenko, 1990). NBT and DAB stained leaf discs and trypan blue leaf staining were quantified using ImageJ software (http://imagej.nih.gov/ij). MDA was measured in tomato leaves according to (Heath and Packer, 1968; Hodges *et al.*, 1999). Suberin staining was done accordingly (Lulai and Morgan, 1992) by applying the leaves 6 hours after wounding with 0.1% neutral red in 0.1M potassium phosphate pH 6.5 for 1 min. Stained leaf area was calculated using Digimizer 3.2.1.0 (http://www.digimizer.com).

The harvested leaves were snap-frozen in liquid nitrogen and stored at −80°C for further use. The first 4 rosette leaves from the bottom were designated as old leaves and the upper-most 4 from top as young leaves.

### Determination of chlorophyll and anthocyanin

For chlorophyll determination, 4 leaf discs were sampled from old, middle and young rosette leaves of WT, *Atxdh1* mutants. The discs (0.7mm diameter) were immersed in 90% EtOH and incubated at 4°C for 2 days in the dark. Absorbance of the extracted chlorophyll was measured at 652 nm and total chlorophyll was estimated (Ritchie, 2006). In addition, the yellow area of the leaf was estimated by employing Digimizer 3.2.1.0 (http://www.digimizer.com) and presented as the ratio of the yellow to total area of leaf, as the chlorophyll damage indicator.

### Determination of Relative Water Content (RWC)

The determination of relative water content was done in sampled detached leaves (8-10th from the bottom) 5 min, 30 min, 6h and 24h after the wounding and fresh weight of the leaf sample was determined. Samples were then placed in de-ionized water in 50 ml falcon tubes to hydrate to high turgidity, at 4 °C for 24 hours. Samples were then removed from water, tightly blotted dry to remove all surface moisture and weighed for turgid weight. Dry weight was determined for samples dried in an oven at 65 °C for 72 hours. The percentage relative water content was measured employing the following formula:

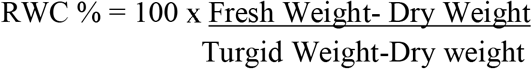

### Membrane cell stability

Osmotic induced electrolyte leakage, as an indicator of membrane damage was determined in five replicates of the same-age attached leaves (middle) after 5min, 30min, 6h and 24h wounding. Five leaves for each replicate were soaked in 10 ml deionized water in falcon tubes and washed using a rotary shaker (500 rpm) for 24 hours at a room temperature to allow solutes to efflux from the tissue. The electrical conductivity of the solution was then measured (initial leakage, Ci). Samples were then autoclaved; the complete leakages of solutes from the autoclaved tissues to the solution were then measured and considered as the maximum electrolyte leakage (Cm). The membrane stability or index of electrolyte leakage was expressed as Ci/Cm × 100 and recorded.

### Metabolites Analysis

Samples (100 mg) were grounded in 25 mM pH 7.5 K_2_PO_4_/KH_2_PO_4_ buffer (1:4 w/v) using a chilled mortar and pestle. The resulting homogenates were transferred to 1.5 ml micro-centrifuge tubes, centrifuged at 15000g for 20 min at 4°C, and the supernatant was used for analyses. Quantification of ureides, allantoin and allantoate, was done according to (Vogels and Van Der Drift, 1970) and glyoxylate was used as blank as described before (Soltabayeva *et al.*, 2018). Xanthine was detected using the xanthine oxidase assay as previously described (Brychkova *et al.*, 2008*a*).

### Protein Extraction and Fractionation

Whole protein from Arabidopsis rosette leaves was extracted as described by (Sagi *et al.*, 1998). Concentrations of total soluble protein in the resulting supernatant were determined according to (Bradford, 1976). Samples for Native/SDS PAGE in gel activity were incubated on ice for 30 min in sample buffer containing 47 mM Tris-HCl (pH 7.5), 2% (w/v) SDS, 7.5% (v/v) glycerol, and 40 mM 1,4-dithio-DL-threitol (DTT) as the thiol-reducing agent, and 0.002% (w/v) bromophenol blue (Sagi and Fluhr, 2001*a*; Srivastava *et al.*, 2017). The incubated samples were centrifuged at 15,000xg for 3 min before loading and subsequently resolved in 7.5% (w/v) SDS-polyacrylamide separating gel and 4% (w/v) stacking gels. Native/SDS PAGE was carried out using 1.5 mm thick slabs loaded with 50 μg of old leaf or 100 μg of young leaf proteins unless otherwise mentioned.

### In-gel XDH activities

Regeneration of the active proteins after denaturing PAGE was carried out by removal of the SDS by shaking the gel for 1 h in 10mM Tris-HCl buffer (pH 7.8) solution (65 ml buffer per ml of gel) containing 2 mM EDTA and 1.0% (w/v) Triton X-100 (Sagi and Fluhr, 2001*b*; Srivastava *et al.*, 2017). Following the regeneration process of the active proteins, the gels were assayed for normal in-gel XDH activities using 0.1mM PMS, 1mM MTT and addition of 0.5mM xanthine mixed with 1mM hypoxanthine in 0.1mM Tris-HCl buffer (pH 8.5), at 25 °C under dark conditions. To detect superoxide generation activity by XDH, 0.25 mM NADH or the mix of xanthine and hypoxanthine was used as specific substrate, respectively, whereas PMS was omitted from the reaction mixture (Yesbergenova *et al.*, 2005). The quantity of the resulting formazan was directly proportional to enzyme activity during a given incubation time, in the presence of excess substrate and tetrazolium salt (Rothe, 1974; Srivastava *et al.*, 2017).

### Western Immunoblotting

Protein crude extract samples (20–50μg) extracted as described by (Sagi *et al.*, 1998), were subjected to Native-PAGE electrophoresis and transferred onto polyvinylidene difluoride membranes (Immun-Blot membranes, Bio-Rad). The membrane was probed with the specific primary antibodies: Anti-AAH (a gift from Claus-Peter Witte, https://www.ipe.uni-hannover.de) at 1:500 dilution ratio, antibody specific to Deg1 protease, autophagy protein 5 (ATG5), chloroplastic Glutamine synthetase (GS2) and pbsO (Agrisera, http://www.agrisera.com) at 1:10,000 ratio to autophagy-related protein 8a (ATG8a) (Abcam, http://www.abcam.com), at 1:1,000 ratio or antibody recognizing large and small subunits of Rubisco (LSU, SSU) (a gift from Prof. Michal Shapira laboratory (http://in.bgu.ac.il/en/natural_science/LifeSciences/Pages/staff/Michal_Shapira.aspx) at a dilution ration of 1:3,000. Thereafter, the proteins underwent binding with secondary antibodies diluted 5000-fold in PBS (anti-rabbit IgG; Sigma-Aldrich). Protein bands were visualized by staining with Clarity Western ECL Substrate (Bio-Rad, USA) and quantified by Image lab (Version 5.2, Bio-Rad, USA).

### Quantitative RT-PCR

Total RNA was extracted from plants using the Aurum Total RNA kit according to the manufacturer’s instructions (Bio-Rad). First-strand cDNA was synthesized in a 10-μl volume containing 350 ng of plant total RNA that was reverse transcribed employing an iScript cDNA Synthesis Kit (Bio-Rad) according to the manufacturer’s instructions. The generated cDNA was diluted 10 times, and quantitative analysis of transcripts was performed employing the set of primers presented in Supplementary Table S1 designed to overlap exon junctions as previously described (Brychkova *et al.*, 2008*a*). Gene expression was normalized to *POLYUBIQUITIN10* (At4g05320) and *EF-1*α (At5g60390) as housekeeping genes, employed as described previously (Soltabayeva *et al*., 2018).

### Statistical analysis

All results are presented as means and standard errors of means. The data for RWC, electrolyte leakage from five independent experiments. Metabolite, protein content and transcripts measurements represent mean obtained through three independent experiments. Each treatment was evaluated using ANOVA (JMP 8.0 software, http://www.jmp.com). Comparisons among three or more groups were made using one-way analysis of variance with Tukey’s multiple comparison tests.

## Results

### Enhanced senescence symptoms in old and middle leaves of *Atxdh1* mutant compared to wound or UV-C treated WT leaves

Wounding and UV-C exerts damaging effects on plants (Reymond *et al.*, 2000; Frohnmeyer and Staiger, 2003; Sharma *et al.*, 2012). Plant recovery was studied by examining fresh weight (FW) of WT and *Atxdh1* mutant leaves 6, 24 and 72 hours after wounding or UV-C irradiation (150mJ/cm2) was applied. The FW of *Atxdh1* mutant increased more slowly than WT and was significantly lower than WT 72 hour later (Supplementary Fig. S3). Wounding resulted in slow FW enhancement in WT, while the weight of *Atxdh1* mutant significantly dropped after 6 hours and then enhanced but was still lower than WT 72 hours after leaf wounding (Supplementary Fig. S3).

The middle leaves subjected to UV-C stress were evaluated for senescence level and revealed higher expression levels of the senescence marker genes, *WRKY53* and *SGN1* (Brychkova *et al.*, 2008*a*), and enhanced yellowing rate of the leaf surface in the middle leaves of *Atxdh1* mutant compared to WT 24h after UV-C treatment (Fig. 1A, C). Notably, 72 hours after application, surface leaf yellowing was noticed in WT and *Atxdh1* mutant middle leaves; however, the rate of yellowing was significantly higher in *Atxdh1* mutant compared to WT (Fig. 1A, B). Interestingly, *Atxdh1* old leaves that were not directly exposed to UV-C, exhibited higher expression levels of senescence genes, as well as a higher chlorophyll degradation rate than those in WT, 72 hours after the UV-C exposure (Fig. 1H, I, J). However young leaves (four leaves from the top) in WT and *Atxdh1* mutant plants were not affected by the UV-C irradiation; differences between genotypes were not detected, being similar to control untreated plants 72 h after the application (Supplementary Fig. S4). Additional, the measurements of MDA, a marker of lipid peroxidation (Khanna-Chopra, 2012; Wang *et al.*, 2013), in middle and old leaves 6, 24 and 72h after wound or UV-C application exhibited higher MDA levels in *Atxdh1* mutant compared to WT at each time point after stress application (Fig. 1G).

**Fig. 1.**
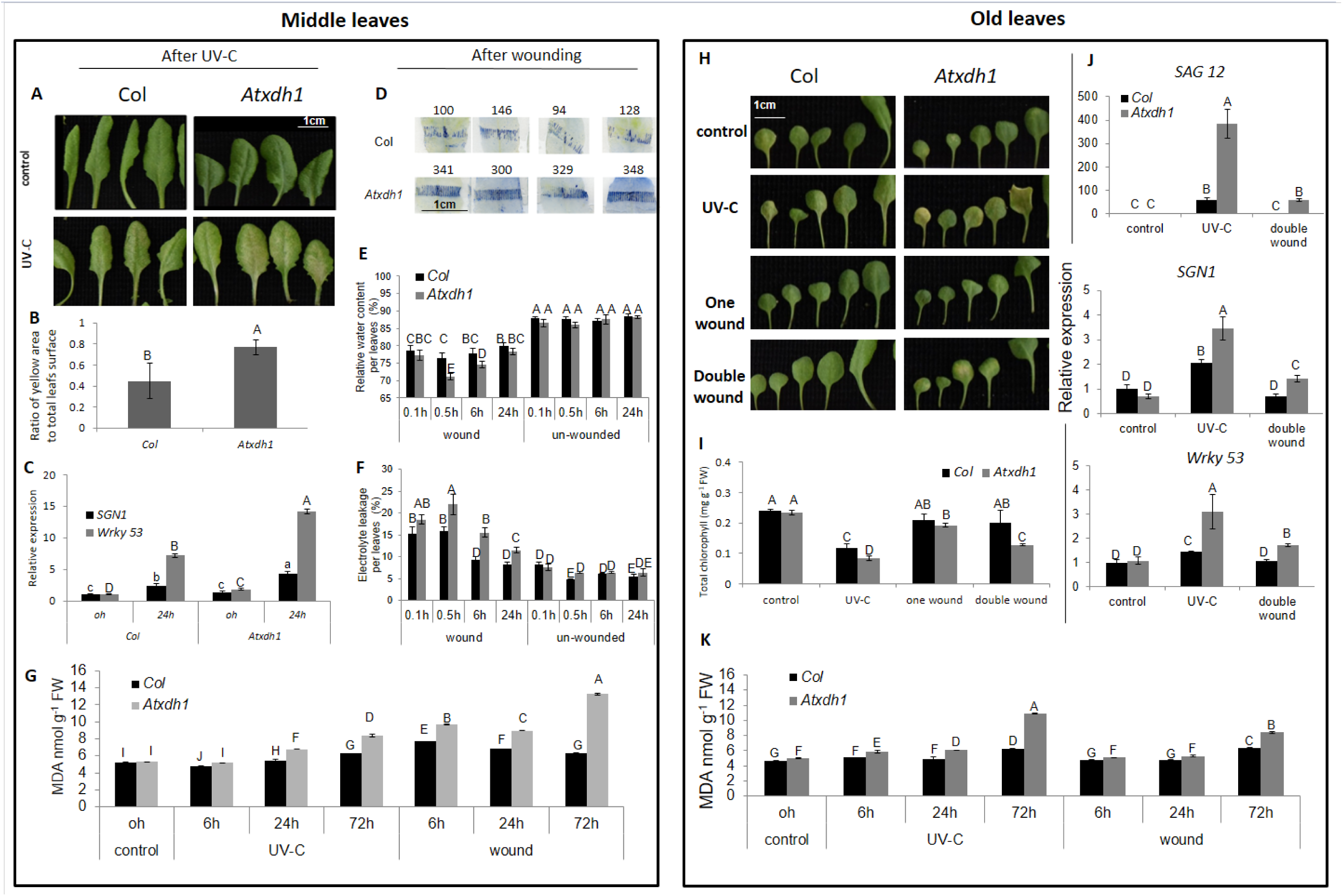
The effect of wounding and UV-C treatments on senescence and damage symptoms in middle and old leaves of WT and *Atxdh1* plants. Left panel (A to G inserts) shows response in middle leaves and right panel (H to K inserts) in old leaves; scale bars in (A), (D) and (H)=1 cm. After UV-C (0.150 J/m2) application the leaves appearance (A) and ratio of yellow area to total leaf surface after 72h (B) and the relative expression of senescence marker *WRKY53* and *SGN1* (C) after 24h of UV-C. Rate of tissue death in wounded area after 24 h (D), the relative water content RWC (E) and electrolyte leakage (F) as affected by wounding in leaves (middle leaves) of WT and *Atxdh1* (SALK 148366). The old leaves appearance (H) and their chlorophyll content (I) after 72 h UV-C, wound and double wound, the relative expression of senescence marker *SAG12*, *WRKY53* and *SGN1* in old leaves (J) after UV-C and wound. The MDA level in middle (G) and old (K) leaves after 6, 24 and 72h of UV-C and wound application of middle leaves compared to 0h, before stress application. The values denoted with different letters are significantly different according to the Tukey-Kramer HSD test; p< 0.05.

The mechanical wounding of middle leaves did not lead to senescence of whole damaged leaves and did not lead to the yellowing of the damaged leaves as UV-C stress did (Supplementary Fig. S5). However, local tissue death was observed in the wounded area of leaves 24h later by trypan blue staining of the middle leaves that was higher in *Atxdh1* leaves compared to WT wounded leaves, reflecting the higher level of cell death in the wound site of mutant plants compared to WT leaves (Fig. 1D). Additionally, the response to wounding was estimated by measuring the relative water content (RWC) at different time points-10min, 0.5, 6 and 24 hours after wounding the middle leaves compared to control non-wounded leaves. The RWC in wounded leaves was significantly decreased as compared to non-wounded in both WT and *Atxdh1* mutant plants at all the time points (Fig. 1E). However, after 0.5 and 6 hours, the RWC in *Atxdh1* mutant leaves was significantly lower than in WT, but 24h after wounding it reached the level of WT wounded leaves (Fig. 1E). This water loss in wounded leaves was consistent with reduction of total leaves weight in *Atxdh1* and WT plants 6 and 24 hour after wounding (Supplementary Fig. S3). Moreover, electrolyte leakage, examined as an indicator of cell membrane integrity, exhibited higher electrolyte leakage in *Atxdh1* mutant leaves than the WT 0.5, 6 and 24 hours after the wounding (Fig. 1F). The higher electrolyte leakage rate and water loss in *Atxdh1* mutant compared to WT wounded leaves indicate the higher rate of cell mortality and damaged membranes in the mutant (Fig. 1 D, F, G), a consequence of plant cell membranes lower healing rate in the wounded leaves. This could also be noticed by the lower FW of *Atxdh1*than WT at later stages of the recovery from wound (Supplementary Fig. S3, right insert). Interestingly, old leaves of *Atxdh1* exposed to double wounding in middle leaves exhibited higher chlorophyll reduction 72h later (Fig. 1H) and enhanced expression of senescence marker genes *SAG12, SGN1* and *WRKY 53* (Fig. 1J) compared to WT old leaves. Yet, young leaves generated new leaves absent of senescence phenotype (Supplementary Fig. S5). Notably, MDA level was enhanced in middle and old leaves in mutant and WT after 72h of UV-C or wounding, but still the MDA level was higher in *Atxdh1* mutant compared to WT (Fig. 1G, K). To this end the results indicate that the middle and old leaves of *Atxdh1* mutant expressed higher sensitivity than WT to UV-C and wounding of middle leaves.

### The higher degradation of chloroplast proteins and their remobilization in untreated, old and *Atxdh1* damaged leaves compared to WT

The chloroplast localized Ribulose-1,5-bisphosphate carboxylase-oxygenase (Rubisco) protein represents ca 50% of the total proteins in mature leaves of C3 plants (Staswick, 1997). Significantly, the enhanced chlorophyll breakdown in old leaves of *Atxdh1* compared to WT after UV-C and/or double wounding (Fig. 1H, I) was parallel with a strong decrease of the large and small subunits of Rubisco [LSU and SSU, respectively (Fig. 2A)]. In middle leaves the LSU and SSU were also reduced in *Atxdh1* mutant compared to WT after UV-C and double wounding (Fig. 2B). Photosystem II oxygen evolving complex (pbsO) protein, a component of PSII (Roose *et al.*, 2010) and Glutamine synthase 2 (GS2), another chloroplast localized protein (Morey *et al.*, 2002) were decreased in old and middle leaves of *Atxdh1* mutant compared to control and WT after UV-C and double wounding of middle leaves (Fig. 2). Notably, the decrease in LSU, SSU, pbsO and GS2 expression by the stress application was stronger in old leaves than in the middle leaves of *Atxdh1*. Additionally, the expression level of the protease Degradation of Periplasmic Proteins 1 (Deg1) involved in the repairment of photosystem II (Kapri-Pardes *et al.*, 2007; Li *et al.*, 2010; Sun *et al.*, 2010), was higher in *Atxdh1* old leaves than WT after double wound or UV-C application, but was not detectable in middle leaves (Fig. 2).

**Fig. 2.**
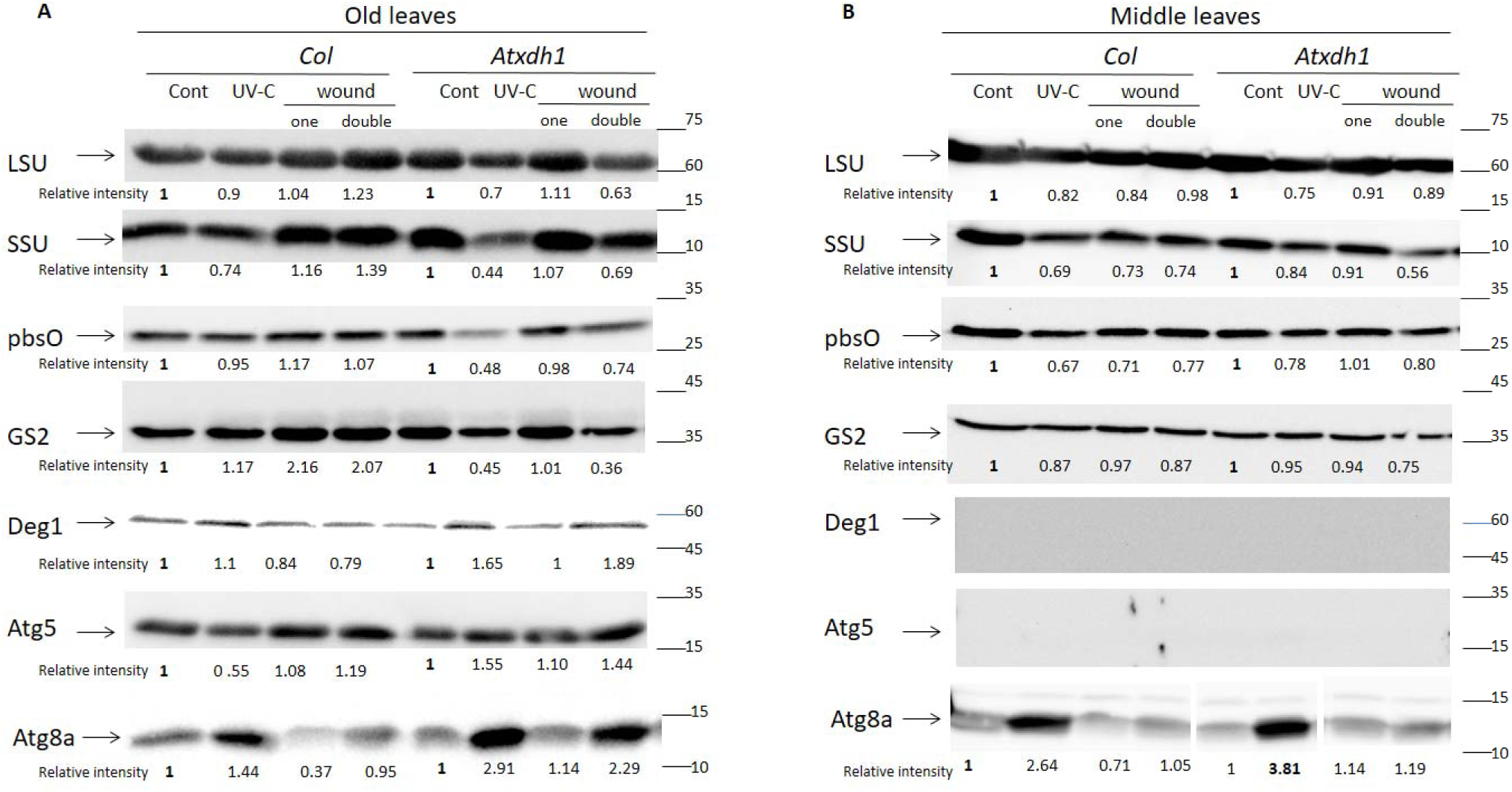
Immunoblot analysis of the large subunit (LSU) and small subunit (SSU) of Rubisco, pbsO proteins, the component of the reaction center of PSII, chloroplastic Glutamine synthetase (GS2), Deg1 protease and autophagy proteins, ATG5 and ATG8a after 72 hours of UV-C, wound, double wound of middle leaves of WT and *Atxdh1*. Proteins were extracted from old (A) leaves (first 4 leaves counted from the bottom) and middle (B) leaves of WT (Col) and *Atxdh1* mutant (SALK_148366) plants, 25 days old. Equal amounts of crude protein extracts were loaded into each lane. The data represent the mean obtained from a representative experiment of two independent biological replications.

The degradation of chloroplast localized proteins and their reutilization involves autophagy related proteins (Ishida and Yoshimoto, 2008). Notably, the analysis of ATG related protein content by immunoblotting employing specific antisera (Yarmolinsky *et al.*, 2014; Soltabayeva *et al*., 2018) revealed increased expression of Autophagy 8a (ATG8a) and Autophagy 5 (ATG5) in *Atxdh1* mutant old leaves than WT after UV-C or double wound were applied to the middle leaves (Fig. 2). Yet, ATG5 was not detectable in middle leaves in WT and *Atxdh1* mutants, whereas ATG8a was higher in *Atxdh1* mutant only after UV-C (Fig. 2), indicating that remobilization of degraded proteins was more active in the old leaves after UV-C irradiation and double wound treatments in *Atxdh1* compared to WT, while in the middle leaves only after UV-C application.

Remarkably, one time wounding of middle leaves didn’t reduce the total chlorophyll content, LSU, SSU, pbsO, GS2 protein content and didn’t increase ATG8a and Deg1 protease in old and middle leaves of *Atxdh1* mutants compared to control or double wounding of *Atxdh1* mutant (Fig. 2), indicating that the senescence of old leaves in the mutant depends on the strength of the stress. Taken together these results indicate that the application of double wound or UV-C in middle leaves results in higher chloroplast protein degradation rate and remobilization than WT in old and middle *Atxhd1* leaves.

### WT leaves exhibited lower ROS level than *Atxdh1* mutant in response to wound or UV-C irradiation

ROS play a role in signaling when they are at low levels (Mittler *et al.*, 2011; Sharma *et al.*, 2012; Baxter *et al.*, 2014) and are known to act as toxic molecules, when generated at higher levels, causing cell death and tissue damage events in plants (Gechev *et al.*, 2006; Van Breusegem and Dat, 2006). It is possible that the higher cell death rate in *Atxdh1* mutant after UV-C or wound application is the result of higher oxidative stress level than in WT leaves. ROS levels were evaluated after UV-C and wound stress in *Atxdh1* and WT plants using nitroblue tetrazolium (NBT) and 3, 3’-diaminobenzidine (DAB) staining to estimate superoxide and hydrogen peroxide level, respectively (Fryer *et al.*, 2002). Twenty four h after UV-C irradiation stress (0.150 J/m^2^) leaf discs were carved from the attached leaves (middle) to carry out DAB or NBT staining. In the local wound treatment, the curved leaf discs were wounded using forceps (Sagi *et al.*, 2004) and 3 hours later were stained with NBT or DAB. UV-C and wounding revealed a higher hydrogen peroxide level in *Atxdh1* mutant as compared to WT (Fig. 3A) and a higher superoxide level in *Atxdh1* only after wounding (Fig. 3). These results are in agreement with the higher MDA level, the results of enhanced lipid peroxidation, in *Atxdh1* mutant than in WT middle leaves (Fig. 1G). Thus, impairment in XDH activity resulted in higher ROS level after wounding and UV-C treatments; likely the result of the lower level of the antioxidant allantoin in the mutant, essential for ROS scavenging as was shown before (Brychkova *et al.*, 2008*a*).

**Fig. 3.**
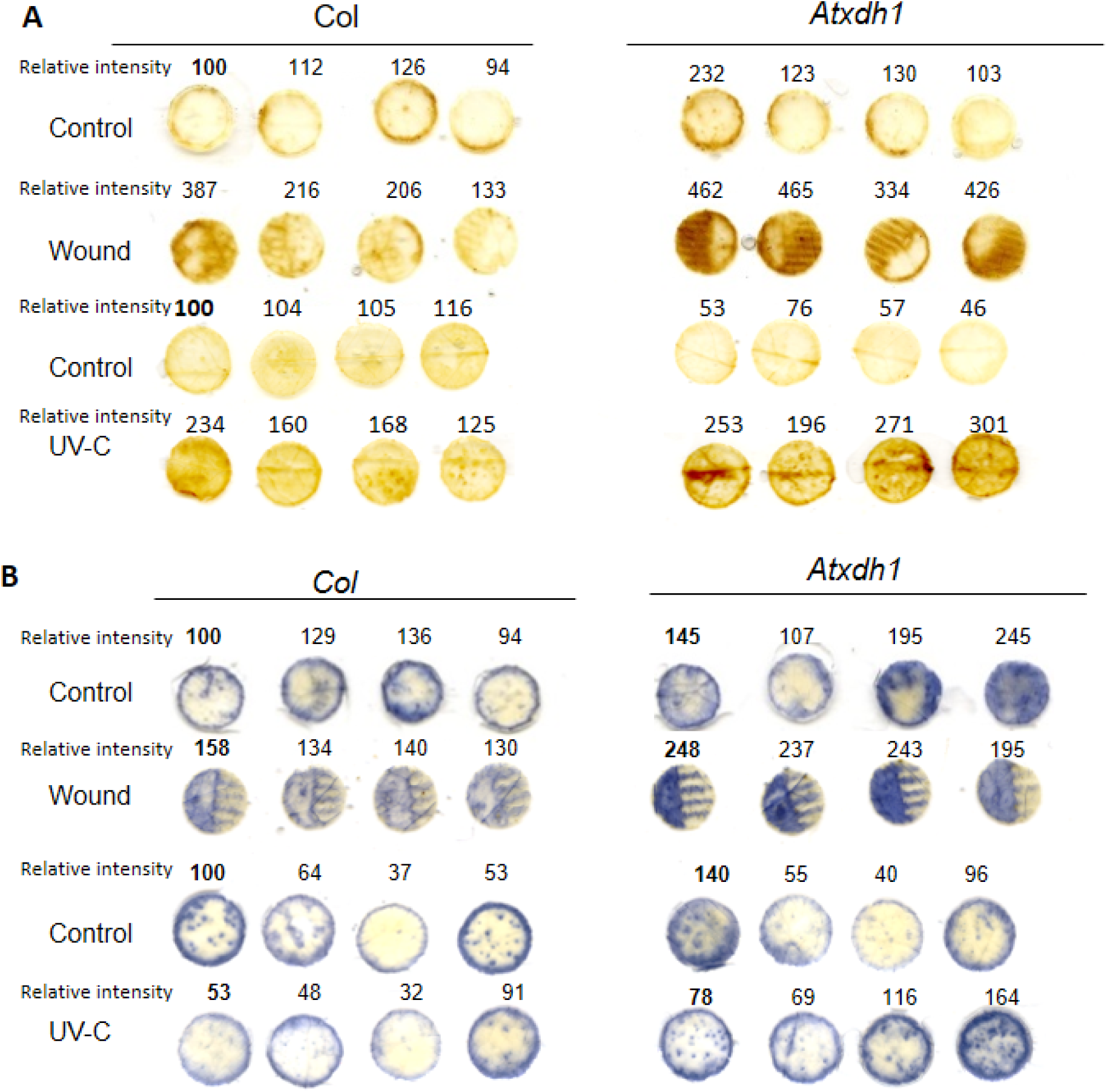
Reactive oxygen species (ROS) accumulation in WT (Col) and *Atxdh1* after UV-C and wounding treatments. Leaf discs sampled from WT (Col) and *Atxdh1* (SALK 148366) leaves after the wound and UV-C irradiation. For hydrogen peroxide detection staining was performed with 3,3’-diaminobenzidine (DAB) (A) and with nitroblue tetrazolium (NBT) for superoxide detection (B) (Brychkova *et al.*, 2008). Wounded leaf discs by forceps were stained for 2 h, 3 hours after wounding and UV-C treated discs were stained 24 h after 150 J/m^2^ UV-C irradiation application. The data represent means obtained from a representative experiment of three independent biological replications.

### Purine degradation was enhanced after wound and UV-C stresses to accumulate purine degraded metabolites in WT and *Atxdh1* mutant

To elucidate the roles of purine degradation in plant responses to UV-C and wounding stresses, the level of up-stream and down-stream purine degraded metabolites was evaluated in WT and *Atxdh1*. The stressed area in middle leaves was compared to the unstressed area of the same rosette leaf as well as to unstressed leaves (Supplementary Fig. S6 A, B). Both stress types; UV-C irradiation as well as wounding, led to a higher accumulation rate of purine metabolites in the damaged zone of leaves as compared to their respective control untreated leaves (Supplementary Fig. S6). Particularly, the xanthine content was strongly accumulated in the damaged area of *Atxdh1* leaves after UV-C and wounding treatment. In WT, xanthine content was also increased however it was lower than in the *Atxdh1* mutant (Supplementary Fig. S6 C, D). The content of the downstream metabolites, allantoin and allantoate, remained generally unchanged in *Atxdh1* mutant leaves following the UV-C or wound, but in WT these metabolites were significantly increased in the damaged leaf area (Supplementary Fig. S6 E-H).

Interestingly, the increase of xanthine content in *Atxdh1* mutant damaged (middle) leaves was observed earlier, 6h after wound application, than UV-C treated middle leaves that was enhanced 24h after the application (Fig. 4 A, B). *Atxdh1* mutant exhibited a higher xanthine accumulation rate after UV-C or wounding not only in middle leaves, but also in old (undamaged) leaves which accumulated more xanthine as compared to the younger leaves, and the increase of xanthine content started later (after 72 h) than the middle leaves (Fig. 4A, B). In WT the xanthine content was also increased more in old leaves compared to young ones, however, the increase was not as prominent as in *Atxdh1* mutants (Fig. 4A, B). The results in the mutant leaves are most likely indicative of a higher xanthine degradation rate in older as well as in the damaged (middle) leaves of WT compared to the youngest leaves.

**Fig. 4.**
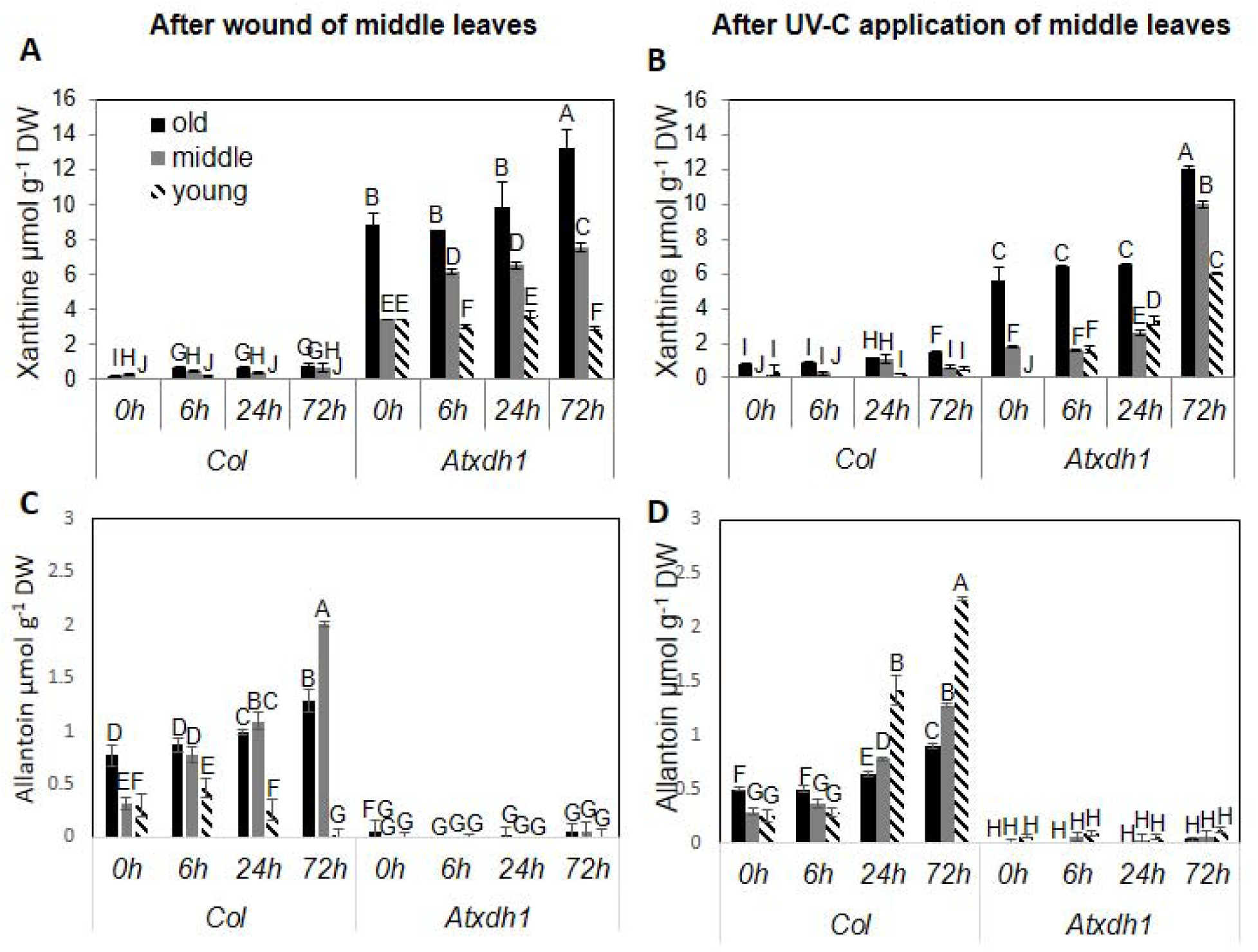
Xanthine (upper panel) and allantoin (lower panel) content changes by time after wound (A and C) and UV-C (B and D) application of middle leaves. 24-day old plants middle leaves exposed to UV-C or wounding and sampled after 6, 24, 72 hours. The 0 hour time point is before treatment application. The data represent means obtained from a representative experiment of three independent biological replications. The values denoted with different letters are significantly different according to the Tukey-Kramer HSD test; p< 0.05.

The generation of the downstream purine catabolism metabolite, allantoin, also started to be enhanced in the middle leaves of WT 6 h after the wounding and 24 h after UV-C application (Fig. 4 C, D). The rise of allantoin content in *Atxdh1* mutant old leaves also started 24 h after the application of UV-C irradiation. In young leaves the allantoin was slightly increased 6 h after the local wound was applied and then reduced, whereas the allantoin level was enhanced 24 h after UV-C irradiation (Fig. 4C, D). Notably, the allantoin was mostly accumulated in middle leaves after wound application, while after UV-C irradiation it was strongly accumulated in young WT leaves (Fig. 4C, D). Notably, the results indicated that after wounding allantoin was remobilized more to damaged leaves, whereas after UV-C treatments allantoin was remobilized to young leaves in WT plants (Fig. 4C, D).

To examine the origin of allantoin mobilization to young and damaged leaves after wounding or UV-C treatments, the middle leaves of WT plants were infiltrated with 1 mM allantoin in the untreated area of wounded or UV-C treated leaves (Supplementary Fig. S7A). The different parts of the tissue were collected from the following three leaf areas: wounded or UV-C treated (marked as W1 or UV1, respectively), untreated areas of leaves (infiltrated area marked as W2 or UV2) and young leaves (marked as young leaves) (Supplementary Fig. S7A). Samples were taken after the infiltration with allantoin, also from water and non-treated plants used as controls (the latter only for the wounding treatment). Six hours after the infiltration a significant increase in allantoin content was evident in the wound or UVC treated area (W1 or UV1) and in young leaves as compared to the infiltrated area [W2 or UV2 (Supplementary Fig. S7A, B and C)]. Interestingly, the allantoin level after 24h was significantly decreased compared to the level detected 6h after allantoin infiltration (Supplementary Fig. S7 B), indicating the higher assimilation rate of the degraded allantoin during the 24 as compared to 6 hours after the application of the wound stress. The enhanced allantoin degradation/consumption was also noticed in WT young leaves after middle leaf wounding (See Fig.4 C), concurrently with a transcript expression increase (See Fig 5 E, G). These results indicate the mobilization of allantoin to the stress treated area and to the young leaves in WT and the activation of ureides degradation in young leaves 24 hours after stress application (recovery period).

**Fig. 5.**
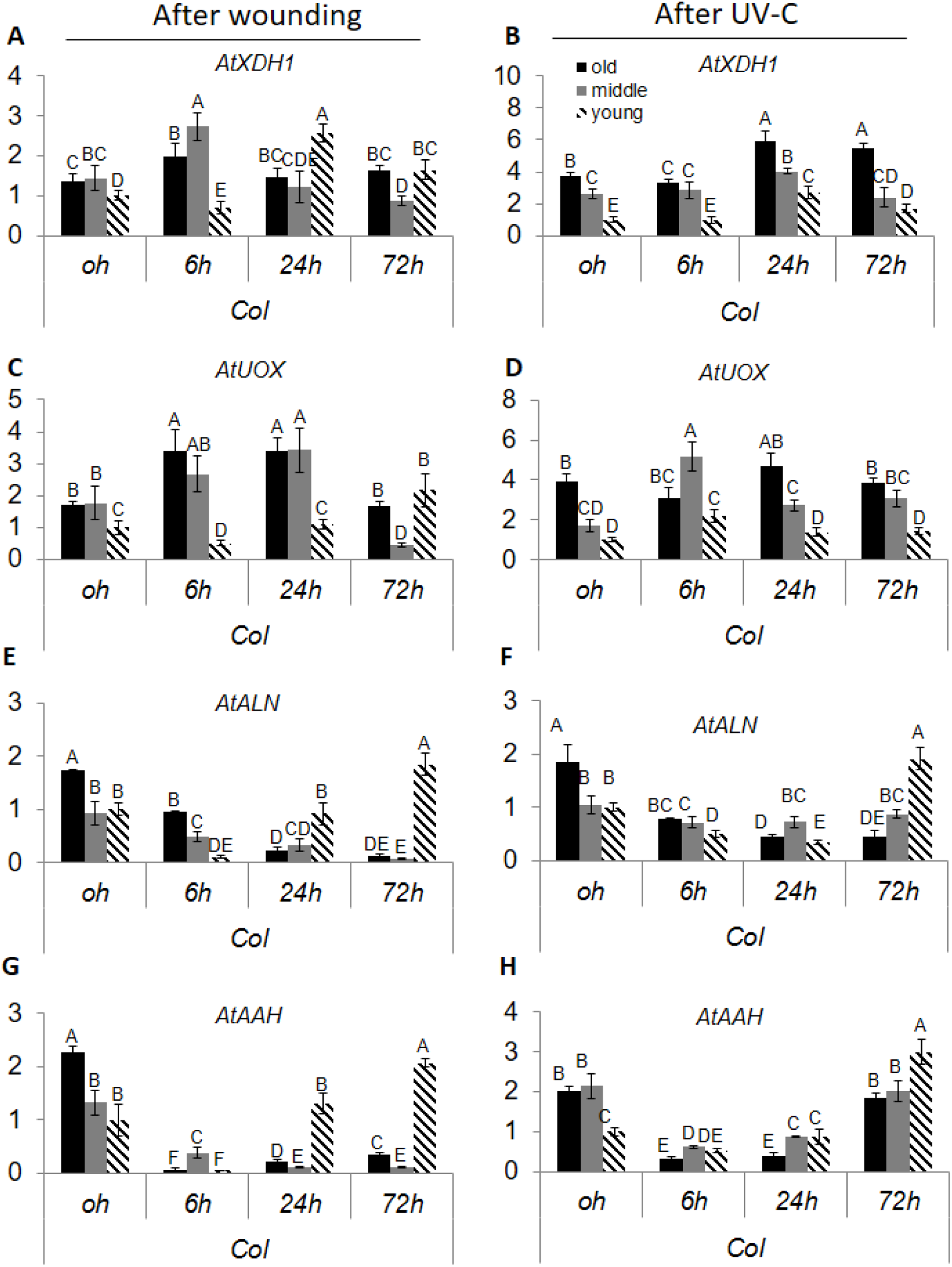
Time course of the transcript expression of the purine catabolism genes after UV-C (right panel) and wounding (left panel) in different leaves of WT. The quantitative analysis of transcripts Xanthine dehydrogenase (*AtXDH*) (A and B), urate oxidase (*AtUOX*) (C and D), allantoinase (*AtALN*) (E and F) and allantoate amidinohydrolase (*AtAAH*) (G and H) was performed by real-time RT-PCR. The expression of genes was compared with the Col at time 0 (before treatment) after normalization to *EF-1*α gene product (*At5g60390*). The data represent means obtained from a representative experiment of three independent biological replications. Values denoted by different letters are significantly different (Tukey-Kramer HSD test, P < 0.05).

### Wound and UV-C stress resulted in similar transcript regulation of purine degradation transcripts

To elucidate whether the accumulation of ureides in WT results from purine catabolism genes network orchestrated by UV-C and wounding stresses, the transcripts of the purine-degrading enzymes were analyzed 0, 6, 24 and 72 h after UV-C and wound application. Xanthine dehydrogenase 1 (*XDH1)* transcript expression level was the lowest in young leaves and was increased in old middle and young leaves 24h after UV-C application (Fig. 5B). In contrast, wounding resulted in *XDH1* transcript increase in old and middle leaves already 6h after stress application but later its expression decreased, while in the young leaves an enhanced expression was noticed 24 after stress application that was still higher than control after 72h (Fig. 5A). The transcript of the urate oxidase (*UOX*), involved in ureides production (Reumann *et al.*, 2007), increased 6h after UV-C application in middle and young leaves and afterwards declined to normal expression in all WT leaves (Fig. 5D). In wound stress, the *UOX* was increased in old and middle leaves of WT after 6h until 72h, but in young leaves it was significantly increased 72h after wound application to the middle leaves (Fig. 5C).

In contrast to the up-stream genes mentioned above, the down-stream genes of the purine degradation pathway, *ALN*, which converts allantoin to allantoate, and *AAH*, which converts allantoate to ureidoglycolate, were decreased after UV-C application in WT up to 24 hours and then *ALN* was strongly increased in young leaves, but not in the other leaves while *AAH* was enhanced in all the leaves especially in the young leaves (Fig. 5F, H). Similarly, the expression of *ALN* and *AAH* was decreased after wounding in old and middle leaves of WT, whereas in young leaves the expression was down regulated up to 6 hours after wounding and later was highly increased in both transcripts (Fig. 5E, G). Thus, transcripts of ureides producers such as XDH and UOX were up-regulated or relatively did not change much, whereas the transcript of ureide degrading enzymes such as *ALN* and *AAH* were down-regulated in both type of stress indicating induction of ureide accumulation as the result of transcript control.

### XDH and AAH proteins expression followed the transcript response after wounding and UV-C stress

To verify the regulation of the purine degradation pathway on protein level in middle leaves after wound or UV-C application, XDH and AAH were chosen as representative proteins of the upstream and downstream components of the purine catabolism and their expression was estimated after UV-C and wounding application. XDH converts xanthine to uric acid and NADH to NAD+ while generating superoxide (Yesbergenova *et al.*, 2005; Zarepour *et al.*, 2010; Werner and Witte, 2011). XDH in-gel activities were estimated by native-SDS PAGE (Yesbergenova *et al.*, 2005). For general XDH in-gel assay the reaction solution contained PMS, MTT and xanthine/ hypoxanthine, while for XDH–NADH in-gel assay that generates superoxide the reaction solution contained MTT and NADH and for the xanthine dependent XDH assay that also generates superoxide the reaction solution contained MTT and xanthine (Yesbergenova *et al.*, 2005). The general XDH activity was increased 6h after the wounding and 72 h after UV-C application (Fig. 6A). The xanthine dependent activity was enhanced after 6h wounding and also after 72 hours of UV-C (Fig. 6B), while the NADH dependent superoxide generating activity of XDH was increased only 24h after the wounding but was not affected by UV-C application (Fig. 6C). This analysis of xanthine or NADH dependent superoxide generating activity of XDH after UV-C or wound stresses demonstrated that the most preferred from the two substrates tested to generate superoxide by XDH was xanthine, likely the consequence of stronger starvation for organic carbon and nitrogen source molecules under the UV-C stress compared to the wound stress.

**Fig. 6.**
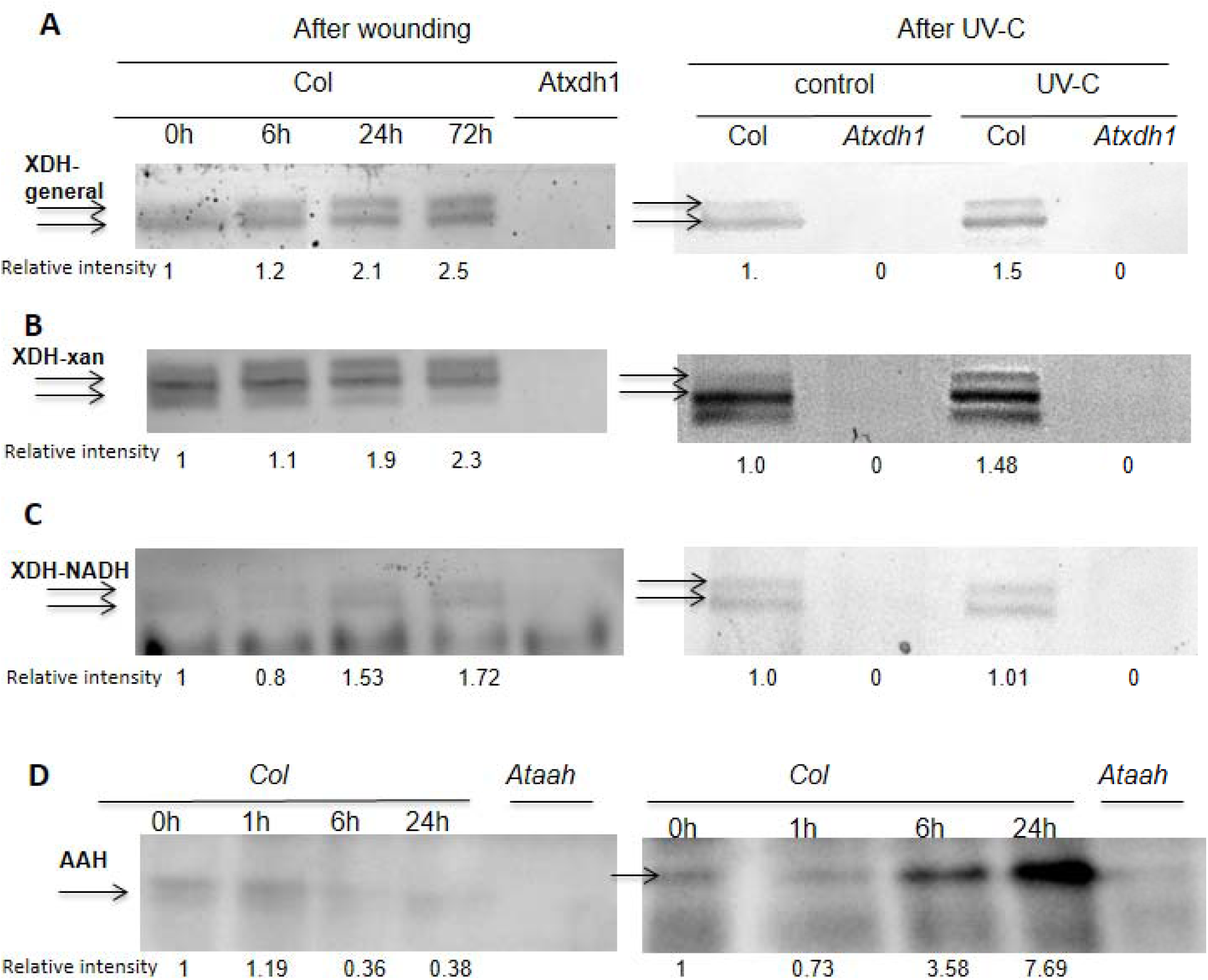
Xanthine dehydrogenase (XDH) activity and Immunoblot analysis of allantoate amidohydrolase (AAH) in middle leaves WT and *Atxdh1* mutant after wounding and UV-C. The xanthine dehydrogenase (A), NADH dependent XDH activity (B) and xanthine dependent XDH activity (C) 6, 24 and 72 hours after wounding and 72 hours after UV-C application. Middle leaves of WT and *Atxdh1* mutant (GABI_049004) were used for the analyses. XDH activities were detected in native-SDS PAGE gel as described in Yesbergenova, et al., (2005). *Atxdh1* (GABI_049004) mutant was used as negative control. Immunoblot analysis of AAH (D) in WT after wounding and UV-C is presented. AAH protein amounts analyzed by Western blot in Native PAGE developed with specific antiserum against AAH and *Ataah* (SALK_112631) mutant was used as negative control. Equal proteins were loaded per lane. The data represent the mean obtained from a representative experiment of two independent biological replications.

AAH protein expression was evaluated by immunoblotting after native PAGE, employing highly specific antisera. The AAH protein was detected as one band in WT and that can be corroborated by its absence in *Ataah* mutant as done before (Soltabayeva *et al*., 2018). The analysis of middle leaves after wounding/UV-C followed the transcript expression showing AAH protein level decrease in WT, 6h after the wounding and in UV-C stress the AAH protein content was decreased 1h after the stress application but was increased 6h after the stress was applied (Fig. 6D). Notably, after 72h, the AAH protein content and native XDH activity were much higher in young and old leaves than in middle leaves after UV-C or wounding (Supplementary Fig. S8). These results indicate that XDH1 activity and AAH protein expression followed the transcript expression after wound and UV-C stresses.

## Discussion

UV-C and wounding have harmful effects on plants (Reymond *et al.*, 2000; Frohnmeyer and Staiger, 2003; Sharma *et al.*, 2012). UV-C irradiation causes DNA damage, oxidation of cellular components (Renzing *et al.*, 1996), and disruption of chloroplasts that results in reduced photosynthesis levels (Booij-James *et al.*, 2000). Wounding results in death of tissue, water loss and may damage the whole plant system (Orozco-Cardenas and Ryan, 1999; León *et al.*, 2001; Reymond *et al.*, 2004). Studies on plant adaptation to UV-C irradiation and wounding stress focus on the role of reactive oxygen species (ROS) regulation (Landry *et al.*, 1995; Conconi *et al.*, 1996; Brosché and Strid, 2003), and on phyto-hormones such as: jasmonic acid (Farmer and Ryan, 1992; Herde *et al.*, 1996), abscisic acid (Pena-Cortes *et al.*, 1989; Herde *et al.*, 1996), ethylene (O’Donnell *et al.*, 1996), and systemin (Pearce *et al.*, 1991). Several reports have shown the role of purine degradation products, allantoin and allantoate, as oxidative stress protectors in plant responses to different stresses (Hasegawa *et al.*, 2000; Brychkova *et al.*, 2008*a*; Alamillo *et al.*, 2010; Kanani *et al.*, 2010; Yobi *et al.*, 2013; Irani *et al.*, 2018). Generally, the increase in levels of allantoin and allantoate was reported under abiotic stresses such as drought (Alamillo *et al.*, 2010; Yobi *et al.*, 2013), salinity (Hasegawa *et al.*, 2000; Kanani *et al.*, 2010; Irani and Todd, 2016; Lescano *et al.*, 2016), cold (Kaplan *et al.*, 2004), extended darkness (Brychkova *et al.*, 2008*a*), high irradiance (Irani *et al.*, 2018) and also in plant response to pathogen attack (Montalbini, 1991). Allantoin and allantoate were shown to act as transportable nitrogen source that can degrade and release the nitrogen under N starvation (Soltabayeva *et al.*, 2018), however the need for the transport of ureides from stress degraded tissues to cope with oxidative stress damage and support growth of young leaves in the recovery period of the stressed plants was hardly shown in non-legume plants.

### Wounding and UV-C stress similarly orchestrated purine degradation pathway to accumulate ureides

Previous studies related to the stress impact on transcripts of purine degradation genes dealing with extended dark stress (Brychkova *et al.*, 2008*a*), salt and drought stresses (Irani and Todd, 2016; Lescano *et al.*, 2016), cadmium toxicity (Nourimand and Todd, 2016) and nitrogen starvation (Soltabayeva *et al.*, 2018), showed two major coordinations of the purine degradation pathway; ureides accumulation or degradation. Yet, it is not clear how it is regulated and how the purine catabolism in the unstressed leaves is affected. Application of UV-C and wound stresses resulted in the accumulation of allantoin in *Arabidopsis* WT as an early response (Fig. 4 C, D). The ureides accumulation was coordinated on transcript level in both types of stresses, where transcripts that are responsible for ureide biosynthesis (such as *XDH1, UOX*) are enhanced, while those responsible for ureide degradation (*ALN* and *AAH*) are reduced, resulting in the accumulation of the ureides (Fig. 5). Further, we also showed that representative proteins upstream and downstream of the purine catabolism pathway, followed the transcripts regulation after UV-C and wounding application (Fig. 6). These results indicate that the accumulation of ureides after wounding or UV-C are controlled on transcript level to result in ureides accumulation. Additionally, in general the later response of the purine degradation pathway to wound or UV-C application (mostly after 72h except for *XDH1* in UV-C treated plants) exhibited a change in transcript response, where the ureide biosynthesis transcripts (*XDH1, UOX*) were declined to the normal expression level, while the ureide degradation transcripts (*ALN* and *AAH*) were increased to normal or higher level (Fig. 5). The AAH and XDH protein expression and activity respectively were enhanced mainly in old and in young leaves 72h after stresses were applied (Supplementary Fig. S8). Interestingly, a similar change in transcripts of purine degradation genes was observed in the recovery stage after extended dark application (Brychkova *et al.*, 2008*a*), suggesting a switching role of ureides from antioxidant ROS scavenger to a metabolite essential for growth as nitrogen and/or carbon source molecule [Supplementary Fig. S2 [Soltabayeva *et al.*, 2018)]. This suggests the dual function of purine catabolism transcriptional control; early response to detoxify by the accumulated ureides the ROS generated by UV-C and wound stresses, as can be seen by the resulting lower hydrogen peroxide and superoxide levels detected in WT compared to *xdh1* mutant leaves (Fig. 3A, B), and a later response of allantoin degradation and remobilization to be used for young leaf growth.

While similar purine degradation coordinated by wound or UV-C stresses was noticed, a different timing of the degraded metabolites’ accumulation in the damaged leaves, differing between the two stresses, was noticed. The changes in purine degraded metabolites in WT and *Atxdh1* were observed 6 h after wounding but only 24 h after the UV-C irradiation (Fig. 4). Yet, 72 h after wounding WT leaves, the ureide degradation genes were still more down regulated compared to the UV-C treated leaves (Fig. 5). These results support the higher ureides level evident in WT wounded leaves compare to UV treated middle leaves (Fig. 4). The results may indicate that the difference in purine degradation rates is associated with the cell death type which differs between the two stresses. In support of this notion is that UV-C causes cell death mostly by apoptotic□like changes (Danon and Gallois, 1998; Gao *et al.*, 2008) while the wound causes necrotic injury (Örvar *et al.*, 1997; Thipyapong and Steffens, 1997).

### Impaired ureides generation enhanced senescence and tissue death in wounded or UV-C treated leaves

UV-C or wound application to middle leaves led to an increase in ROS production in WT and *Atxdh1* mutant leaves. Notably, in *Atxdh1* mutant where ureides generation is blocked as the result of impairment in XDH activity, an enhanced ROS generation compared to WT leaves was demonstrated (Fig. 3). This could be explained by the reduced level of the antioxidant allantoin, which is important in scavenging ROS (Brychkova *et al.*, 2008*a*; Dawood *et al.*, 2021) in *Atxdh1* mutant leaves, resulting in lower antioxidant capacity under the applied stresses. Additionally, allantoin was shown to induce human wound skin healing by increasing collagen deposition (Araújo *et al.*, 2010; Durmus *et al.*, 2012), suggesting that allantoin accumulation after leaf wounding in *Arabidopsis* might heal at least in part, damaged plant cell wall and membranes. Indeed, in support of this notion is the enhanced electrolyte leakage (Fig. 1F) and water loss (Fig. 1G) as well as the lower lignin and suberin formation in *Atxdh1* mutant leaves compared to WT (Supplementary Fig. S9). Additionally, the higher ROS, MDA and tissue death levels (Figs. 1D, G, Fig. 3) in the wounded area of *Atxdh1* mutant leaves compared to WT further support this notion. These results indicate the importance of the purine degradation pathway in response to UV-C and wounding stresses, providing antioxidants to decrease ROS levels and decrease tissue mortality in the damaged leaves and support membrane integrity (see Supplementary Fig. S2).

### Impairment in XDH confers remobilization of degraded proteins from treated and old leaves in response to wound or UV-C irradiation

Similar to the treated leaves, *Atxdh1* mutant old leaves also showed accelerated senescence 72 hours after UV-C and double wounding application. This was followed by a higher chlorophyll degradation rate (Fig. 1 H, I) and more decreased chloroplast protein levels (Fig. 2A, B) than WT. Notably, levels of proteins that function in degraded proteins remobilization, the Atg8 and Atg5 (Klionsky, 2007; Mizushima, 2007; Xie and Klionsky, 2007), were higher in the old leaves of *Atxdh1* mutant compared to WT (Fig. 2A). Since Atg8 and Atg5 are activated during carbon or nitrogen starvation (Doelling *et al.*, 2002; Xie and Klionsky, 2007; Chung *et al.*, 2009; Guiboileau *et al.*, 2012; Honig *et al.*, 2012) these results suggest that the senescence in old leaves of *Atxdh1* mutant are the results of a shortage in internal nutrients, essential to be remobilized to growing leaves and/or to reproductive or storage organs (Vierstra, 1996; Hopkins *et al.*, 2007; Pyung *et al.*, 2007). In contrast, in WT old leaves, a lower protein degradation rate was observed [See LSU, SSU, pbsO, GS2 and Deg1chloroplast localized proteins (Fig. 2A)], likely the result of the enhanced purine degradation rate, noticed on transcript (Fig. 5) and protein levels (Fig. 6), as well as purine degraded metabolites level (Fig. 4C, D) after UV-C or wound application to the middle leaves. Notably, increasing the local wound area from one wound to double wound resulted in enhanced senescence symptoms in *Atxdh1* mutant old leaves as compared to WT, 72h after wound application (Fig. 1H, I, J). Accordingly, in old leaves of WT the generation of ureides was increased with the increase of wound area (double wound) in the middle leaves (Fig. S9), whereas in *Atxdh1* the chloroplast proteins were degraded more than in WT and were remobilized from the old leaves as indicated by Atg5 and Atg8a enhanced protein expression (Fig 2 A). These results indicate the essentiality of ureides generation in old leaves to cope with stress and its consequences, avoiding protein degradation for nutrient supply at the shortage of purine degraded product such as in *Atxdh1* mutants. Additionally, the results indicate that ureide level in old leaves depends on stress intensity.

### Purine degradation level in middle leaves positively responds to the intensity of UVC/wound stresses

The damage level in middle leaves differed between UV-C and wound stress. UV-C irradiation resulted in a whole surface of yellowing that was larger in *Atxdh1* than in wild-type (Fig. 1A), while after local wounding, the damaged area (cell death) was limited to the wounded area (Fig. 1D), whereas the un-wounded area of leaves didn’t show any yellowish coloring (Supplementary Fig. S5). Additionally, UV-C treatment, but not wound damage, induced protein remobilization in the middle leaves as can be seen by Atg8 enhanced expression, (Fig. 2B). This suggests that leaves damaged by UV-C irradiation were a source of macronutrients for growth of new young leaves. In support of this notion, allantoin accumulation in young leaves, as well as in the middle-treated leaves (albeit at lower levels than in the young leaves) after UV-C irradiation (Fig. 4), suggests that the accumulated ureides were remobilized from treated leaves to young leaves in WT. Further support of this notion was revealed by subjecting *Ataln* mutant, impaired in degrading allantoin to allantoate, to UV-C irradiation. An enhanced accumulation rate of allantoin in middle treated leaves and a much higher increase in the young leaves was evident after UV-C treatment. Wounding *Ataln* mutant resulted in allantoin accumulation in both treated middle leaves as well as young leaves (Supplementary Fig. S10). Additionally, the increase of damage level by enhancing the wounded zone led to an increase of ureides level in the damaged and old leaves of WT and in *Ataln* mutants compared to *Atxdh1* mutant (Supplementary Fig. S10). These results indicate that the purine degradation level in middle leaves positively responds to the intensity level of UVC or wound stresses.

### Ureides support plant recovery from wound as antioxidant, while under UV-C, as nutrient for young leaves growth

Ureides, are known to function as major root to shoot nitrogen transport compounds in legumes (Schubert, 1986; Amarante *et al.*, 2006). Active mobilization of ureides as an internal nitrogen source from old to young leaves was shown in *Arabidopsis* plants exposed to nitrogen starvation (Soltabayeva *et al.*, 2018). Yet, ureides were also shown to act as an efficient antioxidants in plants exposed to extended dark stress (Brychkova *et al.,* 2008). Remarkably, compared to the old leaves, 72h after stress application allantoin was strongly accumulated in middle leaves in response to wounding and in young leaves after UV-C irradiation (Fig. 4C, D). These results suggest that purines from old leaves in WT remobilize to young or middle leaves during the recovery period, either as a nitrogen source molecule and/or to protect leaves tissue from oxidative stress, acting as an antioxidant molecule as shown before (Brychkova *et al.*, 2008) (Supplementary Fig. S2, Fig. 7).

**Fig. 7.**
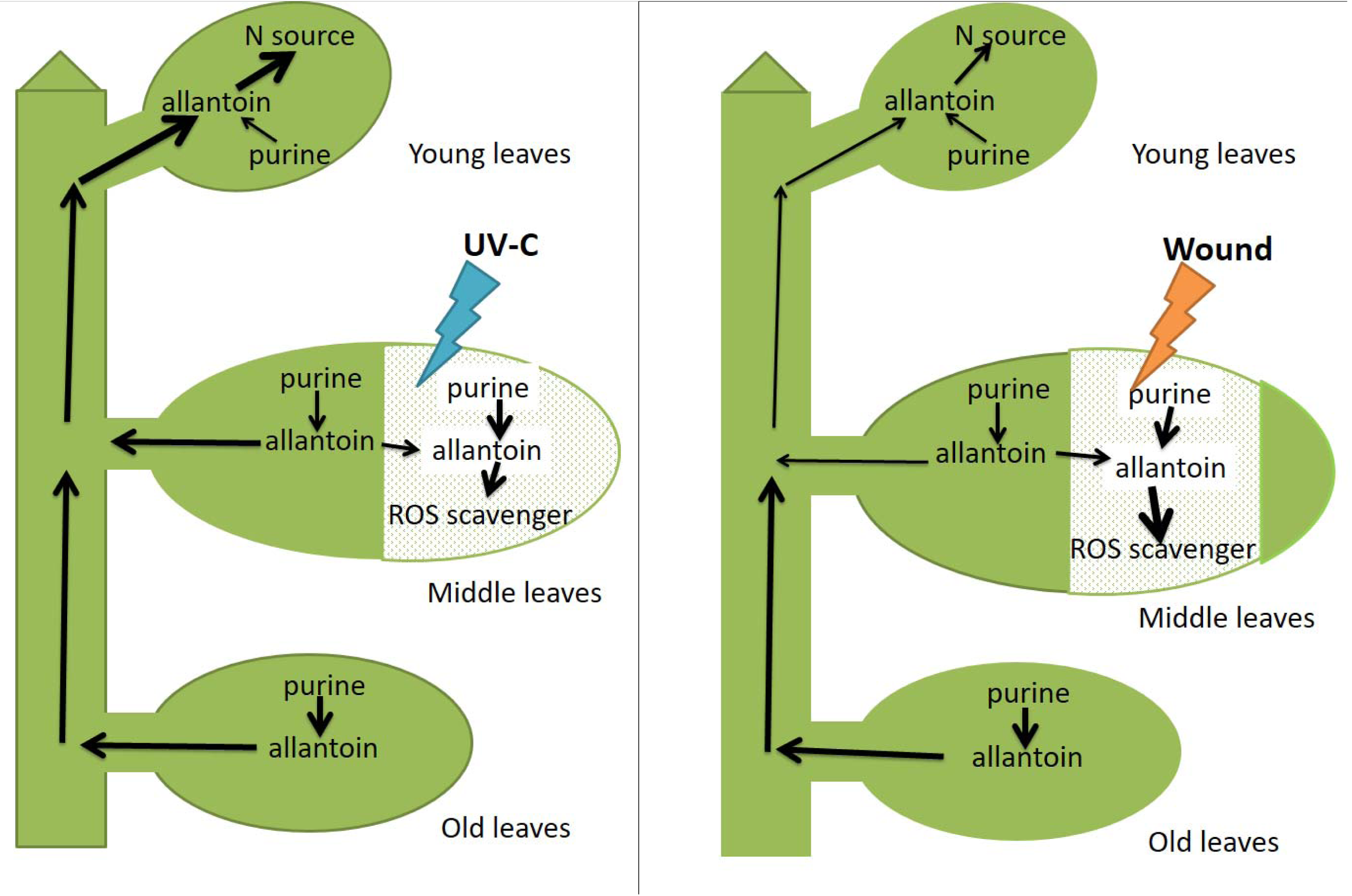
Schemes describing the degradation of purines and the remobilization of the degraded product, allantoin in *Arabidopsis* WT leaves in response to UV-C (left insert) or wound (right inserts) stresses applied to the middle leaves. The middle leaves (5^th^ to 10^th^ counted from the bottom) were treated with UV-C irradiation (0.150 J/m2) or wounding (with forceps). Damaged areas in leaves are shown as a white dotted area. Arrow indicates the direction of remobilized allantoin pool (degraded purine product) and enhanced thickness of an arrow indicates enhanced level (or rate) of the remobilized allantoin. Remobilization of allantoin differs between leaves. Treating the middle leaves with UV-C or wounding resulted in enhanced supply of allantoin, the purine degradation product, in old leaves and in its remobilizing to the young leaves likely acting as N source as shown in Soltabayeva *et al*., (2018). In the middle leaves, the generated allantoin strongly accumulated within the middle leaves, especially from the damaged leaves area but from the undamaged side of leaves as well, (see arrow thickness), likely acting as ROS scavenger as shown in Brychkova *et al*., (2008). The allantoin originated from the middle leaves was remobilized not only to the damaged area but may move to the young leaves. Later, during the recovery period after 72h, allantoin accumulated after UV-C irradiation was strongly remobilized to the young leaves from the old and the middle-damaged leaves at a higher level than the mobilization of allantoin in case of wound stress.

Notably, AAH, the allantoin degrading enzyme, was higher in young than the middle (treated) leaves 72 h after UV-C and wound application, indicating a higher rate of allantoin degradation in young leaves compared to damaged leaves in treated plants, as well as in treated leaves compared to the control untreated middle leaves (Supplementary Fig. S8 B). These results, supported by the infiltration of 1 mM allantoin to the untreated area of middle leaves subjected to the UV-C and wound stresses (see Supplementary Figs. S7 and S9), indicate that after UV-C application, purines remobilize in WT from damaged and old leaves to young leaves to provide nitrogen and/or antioxidant molecules. In contrast, after wounding the ureides are remobilized to the young leaves, yet more to the treated leaves, functioning as antioxidant and/or healing agents in the middle wounded leaves (see scheme in Fig. 7).

### Summary

In the current study it was shown that ureides are similarly accumulated in response to UV-C irradiation and wound but differently remobilized during recovery in *Arabidopsis* leaves. The UV-C and wound stresses coordinated a purine degradation pathway in similar pattern to accumulate the ureides in the damaged zone to scavenge the generated ROS (Fig. 3, Supplementary Fig. S6, Fig. 4 and Fig. 7). This accumulation of ureides triggered by UV-C and wounding was mainly in the damaged area as well as in the oldest leaves (Fig. 4, Fig. 7). Later, during recovery period, the ureides accumulated in old and middle leaves were remobilized to damaged and/or young leaves depending on the stress type (Fig.4 and Fig. S5-6, Fig. S9, Fig. 7). After wounding they were mainly concentrated in damaged leaves and after UV-C were strongly remobilized to young leaves (Fig. 4 and Fig. S9, scheme in Fig. 7). The remobilized ureides were actively degraded in young leaves and/ or damaged leaves to provide nitrogen source (Fig. 5–6, Fig. S7 and Fig. S9).

## Supplementary data

Fig. S1. Microarray analysis of the “purine network” transcripts in response to selected stressors.

Fig. S2. Scheme describing the orchestration of purine degradation pathway transcripts by different stresses resulting in either ROS scavenging by the accumulated ureides or providing N source, the result of ureides degradation.

Fig. S3. Time course of the effect of wounding and UV-C treatments on shoot fresh weight of WT and *Atxdh1* plants.

Fig. S4. The leaves’ appearance (A) and total chlorophyll content (B) of WT and *Atxdh1* plants after 72 hours of UV-C (0.150 J/m2) irradiation.

Fig. S5. The appearance of WT and *Atxdh1* plants leaves 72 hours after UV-C (0.150 J/m2) irradiation or wound, compared to leaves of control untreated plants.

Fig. S6. Xanthine, allantoin, and allantoate content in response to UV-C stress (A) and wounding (B) in damaged and undamaged areas of WT and *Atxdh1* treated leaves.

Fig. S7. Allantoin content in wound or UV-C treated, untreated and young leaves of WT after infiltration with 1mM allantoin.

Fig. S8. Xanthine dehydrogenase (XDH) activity and Immunoblot analysis of allantoate amidohydrolase (AAH) in all leaves of WT and *Atxdh1* mutant 72h after wounding or UV-C irradiation.

Fig. S9. Suberin staining in the wounded area of WT and *Atxdh1* (SALK 148366) mutant leaves.

Fig. S10. Allantoin content after UV-C, one and two forceps wound application to WT, *Atxdh1* (SALK 148366) and *Ataln* (SALK_ 013427) middle leaves.

Table S1. Gene-specific sequences of primers used for the expression analyses

## Acknowledgements

This research was supported by the Israel Center of Research Excellence (ICORE) “Plant Adaptation” (ISF Grant no. 757/12) and the Gerda Frieberg Chair in Agriculture Water Management.

## Author Contribution

A.S. participated in designing the research plans and performed the experiments and analyses. S.S participated in XDH in gel assay; A.K. participated in DAB staining; D.O and Z. N participated in Immunoblot analysis, D.S in ureides content analysis, A.B. participated in qRT-PCR. M.S. conceived the original idea, designed the research plan, and supervised the research work. The manuscript was jointly written by A.S. and M.S.

## Data availability statement

All data supporting the findings of this study are available within the paper and within its supplementary materials published online.

Arabidopsis sequence data from this article can be found on TAIR (https://www.arabidopsis.org/) under accession numbers: AAH, At4g20070; ALN, At4g04955; *UOX*, *At2g26230*; *XDH1, At4g34890*; *EF-1*α*, At5g60390, UBQ10, At4g05320*; *WRKY53, At4g23810; SGN1, At4G22920; SAG12, AT5G45890;* ATG5, AT3G51830; GS2, AT5G35630; pbsO, AT5G66570; Deg1, AT3G27925; ATG8a, AT4G21980; LSU, ATCG00490; SSU, AT1G67090.

