## Supplementary material for "Ureides are similarly accumulated in response to UV-C irradiation and wound but differently remobilized during recovery in *Arabidopsis* leaves": Supp. Figures 1-10 and Table 1

### Slide 1
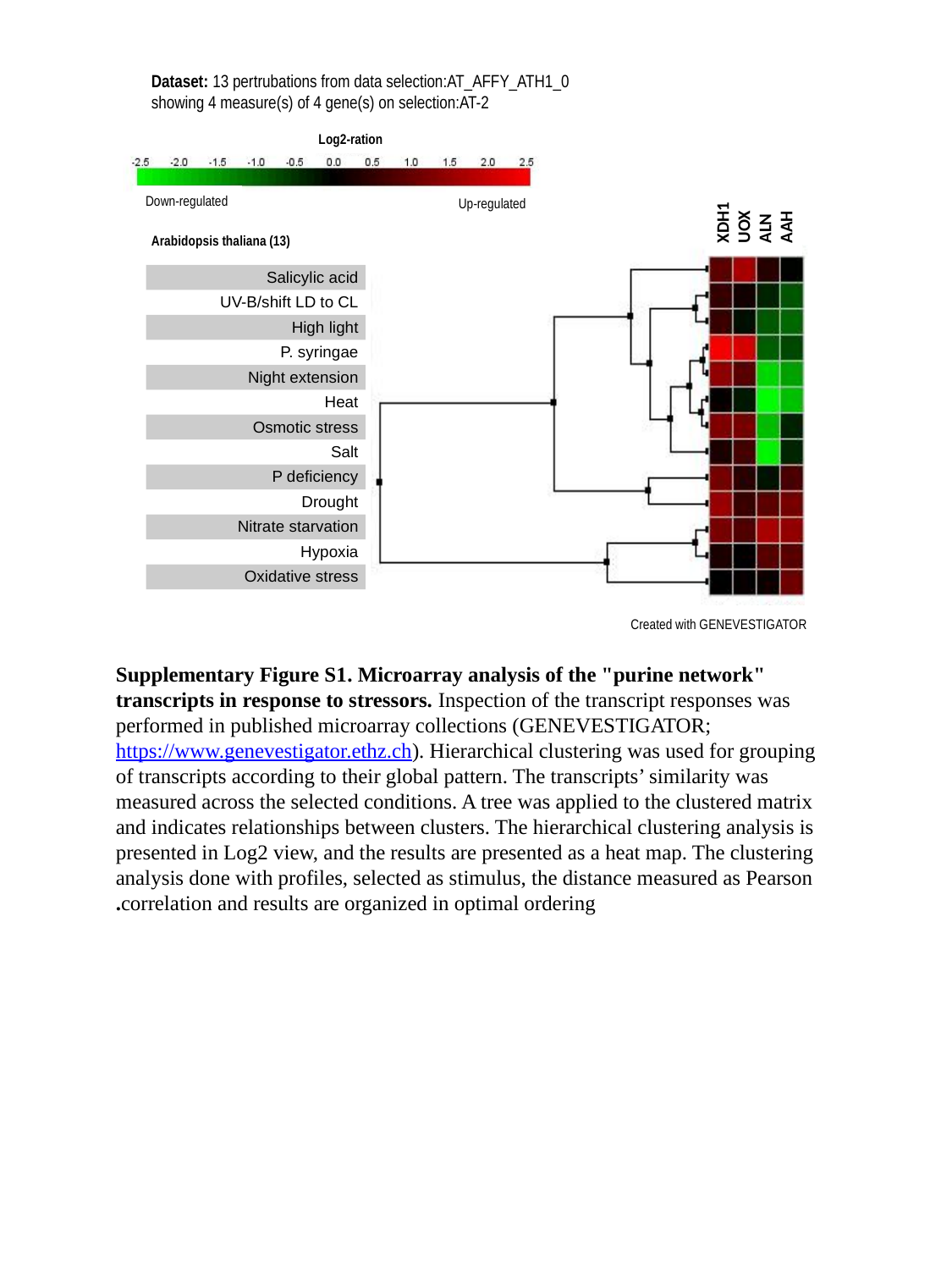

Dataset: 13 pertrubations from data selection:AT_AFFY_ATH1_0
 showing 4 measure(s) of 4 gene(s) on selection:AT-2
Log2-ration
XDH1
UOX
ALN
AAH
Down-regulated
Up-regulated
Arabidopsis thaliana (13)
| Salicylic acid |
| --- |
| UV-B/shift LD to CL |
| High light |
| P. syringae |
| Night extension |
| Heat |
| Osmotic stress |
| Salt |
| P deficiency |
| Drought |
| Nitrate starvation |
| Hypoxia |
| Oxidative stress |
Created with GENEVESTIGATOR
Supplementary Figure S1. Microarray analysis of the "purine network" transcripts in response to stressors. Inspection of the transcript responses was performed in published microarray collections (GENEVESTIGATOR; https://www.genevestigator.ethz.ch). Hierarchical clustering was used for grouping of transcripts according to their global pattern. The transcripts’ similarity was measured across the selected conditions. A tree was applied to the clustered matrix and indicates relationships between clusters. The hierarchical clustering analysis is presented in Log2 view, and the results are presented as a heat map. The clustering analysis done with profiles, selected as stimulus, the distance measured as Pearson correlation and results are organized in optimal ordering.

### Slide 2
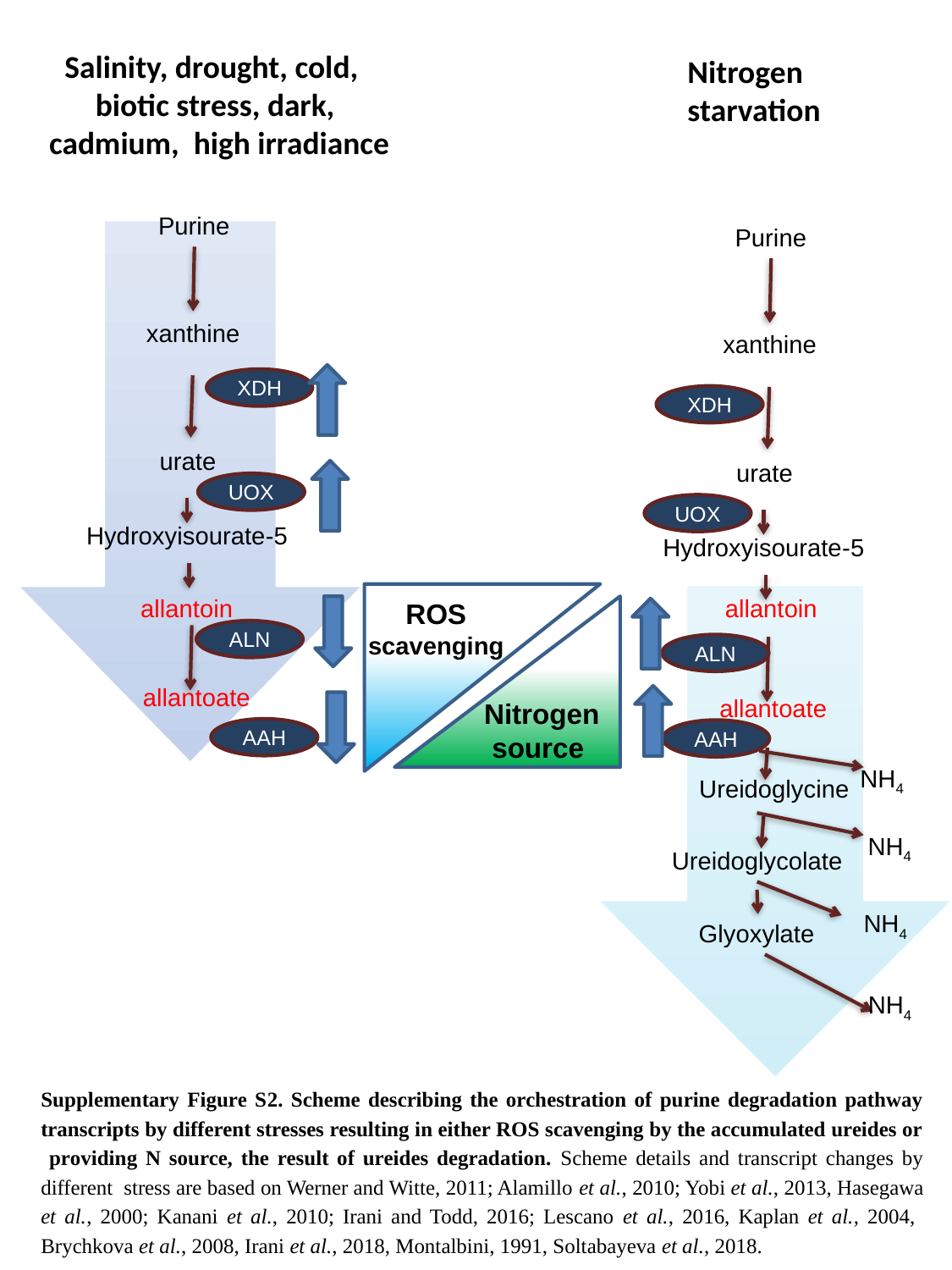

Salinity, drought, cold, biotic stress, dark, cadmium, high irradiance
Nitrogen starvation
Purine
xanthine
XDH
urate
UOX
5-Hydroxyisourate
allantoin
ALN
allantoate
AAH
Purine
xanthine
XDH
urate
UOX
5-Hydroxyisourate
allantoin
ALN
allantoate
AAH
NH_4_
Ureidoglycine
NH_4_
Ureidoglycolate
NH_4_
Glyoxylate
NH_4_
ROS scavenging
Nitrogen
 source
Supplementary Figure S2. Scheme describing the orchestration of purine degradation pathway transcripts by different stresses resulting in either ROS scavenging by the accumulated ureides or providing N source, the result of ureides degradation. Scheme details and transcript changes by different stress are based on Werner and Witte, 2011; Alamillo et al., 2010; Yobi et al., 2013, Hasegawa et al., 2000; Kanani et al., 2010; Irani and Todd, 2016; Lescano et al., 2016, Kaplan et al., 2004, Brychkova et al., 2008, Irani et al., 2018, Montalbini, 1991, Soltabayeva et al., 2018.

### Slide 3
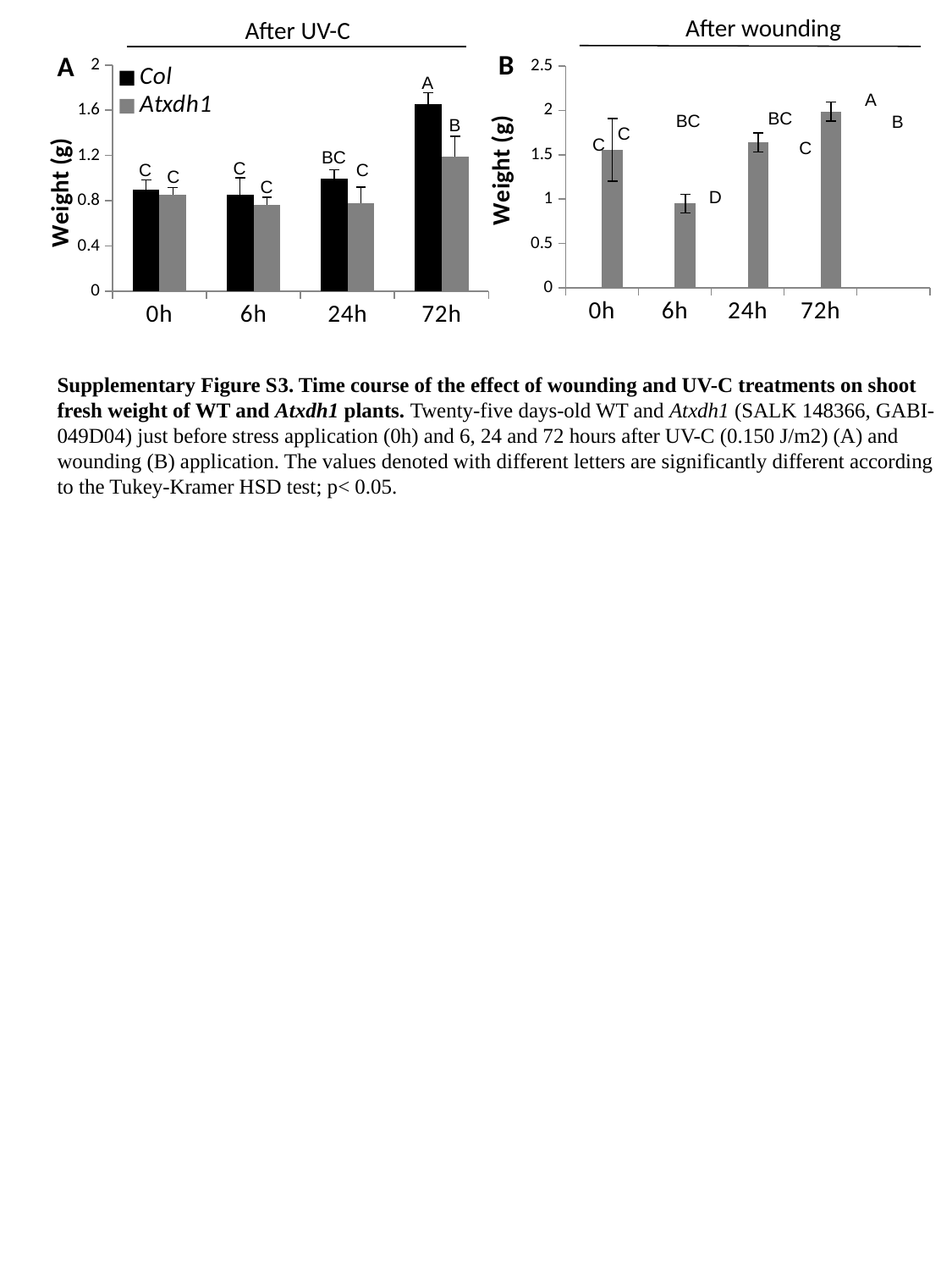

After wounding
After UV-C
#### Chart
| Category | Col | Atxdh1 |
|---|---|---|
| 0h | 0.8999999999999999 | 0.8545 |
| 6h | 0.8506499999999999 | 0.761000000000001 |
| 24h | 0.9984999999999999 | 0.778 |
| 72h | 1.654 | 1.1875 |
#### Chart
| Category | Col | Atxdh1 |
|---|---|---|
| 0h | 1.61 | 1.555 |
| 6h | 2.01573333333333 | 0.9510000000000001 |
| 24h | 2.019 | 1.64 |
| 72h | 2.353 | 1.988 |B
A
A
A
BC
BC
B
B
C
C
C
BC
C
C
C
C
C
D
Supplementary Figure S3. Time course of the effect of wounding and UV-C treatments on shoot fresh weight of WT and Atxdh1 plants. Twenty-five days-old WT and Atxdh1 (SALK 148366, GABI-049D04) just before stress application (0h) and 6, 24 and 72 hours after UV-C (0.150 J/m2) (A) and wounding (B) application. The values denoted with different letters are significantly different according to the Tukey-Kramer HSD test; p< 0.05.

### Slide 4
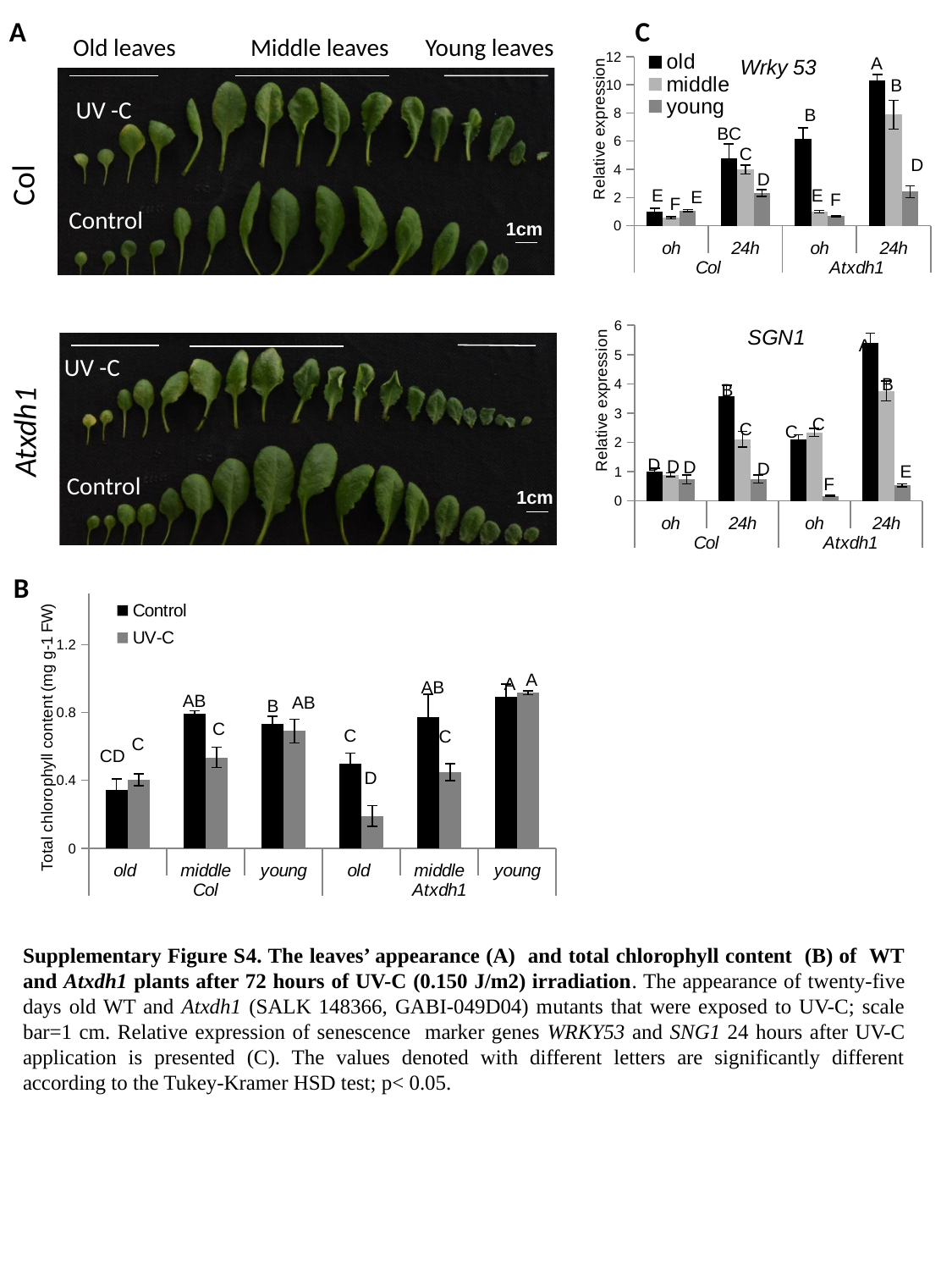

A
C
Old leaves
Middle leaves
Young leaves
#### Chart: Wrky 53
| Category | old | middle | young |
|---|---|---|---|
| oh | 1.0 | 0.554480221060753 | 1.062884633106392 |
| 24h | 4.769109260285834 | 3.9756503616577765 | 2.3106501981226573 |
| oh | 6.1634991157904615 | 0.9923188544811221 | 0.6728986187639224 |
| 24h | 10.309307037111342 | 7.882724418552347 | 2.405348295729361 |A
B
UV -C
B
BC
C
D
Col
D
E
E
E
F
F
Control
1cm
#### Chart: SGN1
| Category | old | middle | young |
|---|---|---|---|
| oh | 1.0 | 0.8881181460287135 | 0.7312292919159598 |
| 24h | 3.5672429569915405 | 2.110049975992757 | 0.7501279910699842 |
| oh | 2.108073678065253 | 2.337656276672854 | 0.17050136738673885 |
| 24h | 5.391113166292409 | 3.7525929006549923 | 0.5322424603861655 |A
UV -C
B
B
C
Atxdh1
C
C
D
D
D
D
E
Control
F
1cm
B
#### Chart
| Category | Control | UV-C |
|---|---|---|
| old | 0.34450000000000003 | 0.4025 |
| middle | 0.7954999999999999 | 0.5359999999999999 |
| young | 0.7324999999999999 | 0.691 |
| old | 0.501 | 0.19 |
| middle | 0.7725 | 0.448 |
| young | 0.8925 | 0.9175 |A
A
AB
AB
AB
B
C
C
C
C
CD
D
Supplementary Figure S4. The leaves’ appearance (A) and total chlorophyll content (B) of WT and Atxdh1 plants after 72 hours of UV-C (0.150 J/m2) irradiation. The appearance of twenty-five days old WT and Atxdh1 (SALK 148366, GABI-049D04) mutants that were exposed to UV-C; scale bar=1 cm. Relative expression of senescence marker genes WRKY53 and SNG1 24 hours after UV-C application is presented (C). The values denoted with different letters are significantly different according to the Tukey-Kramer HSD test; p< 0.05.

### Slide 5
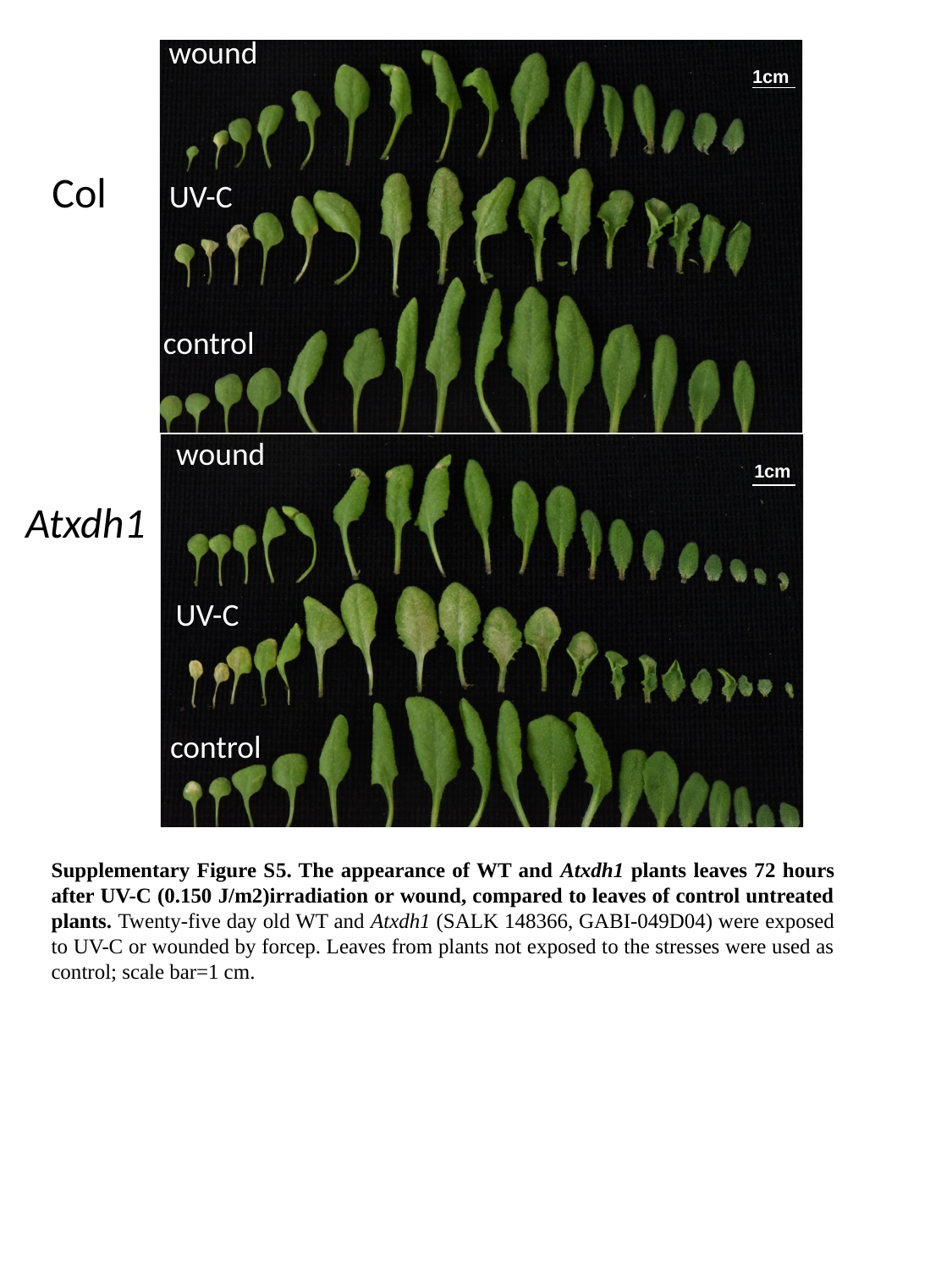

wound
1cm
Col
UV-C
control
wound
1cm
Atxdh1
UV-C
control
Supplementary Figure S5. The appearance of WT and Atxdh1 plants leaves 72 hours after UV-C (0.150 J/m2)irradiation or wound, compared to leaves of control untreated plants. Twenty-five day old WT and Atxdh1 (SALK 148366, GABI-049D04) were exposed to UV-C or wounded by forcep. Leaves from plants not exposed to the stresses were used as control; scale bar=1 cm.

### Slide 6
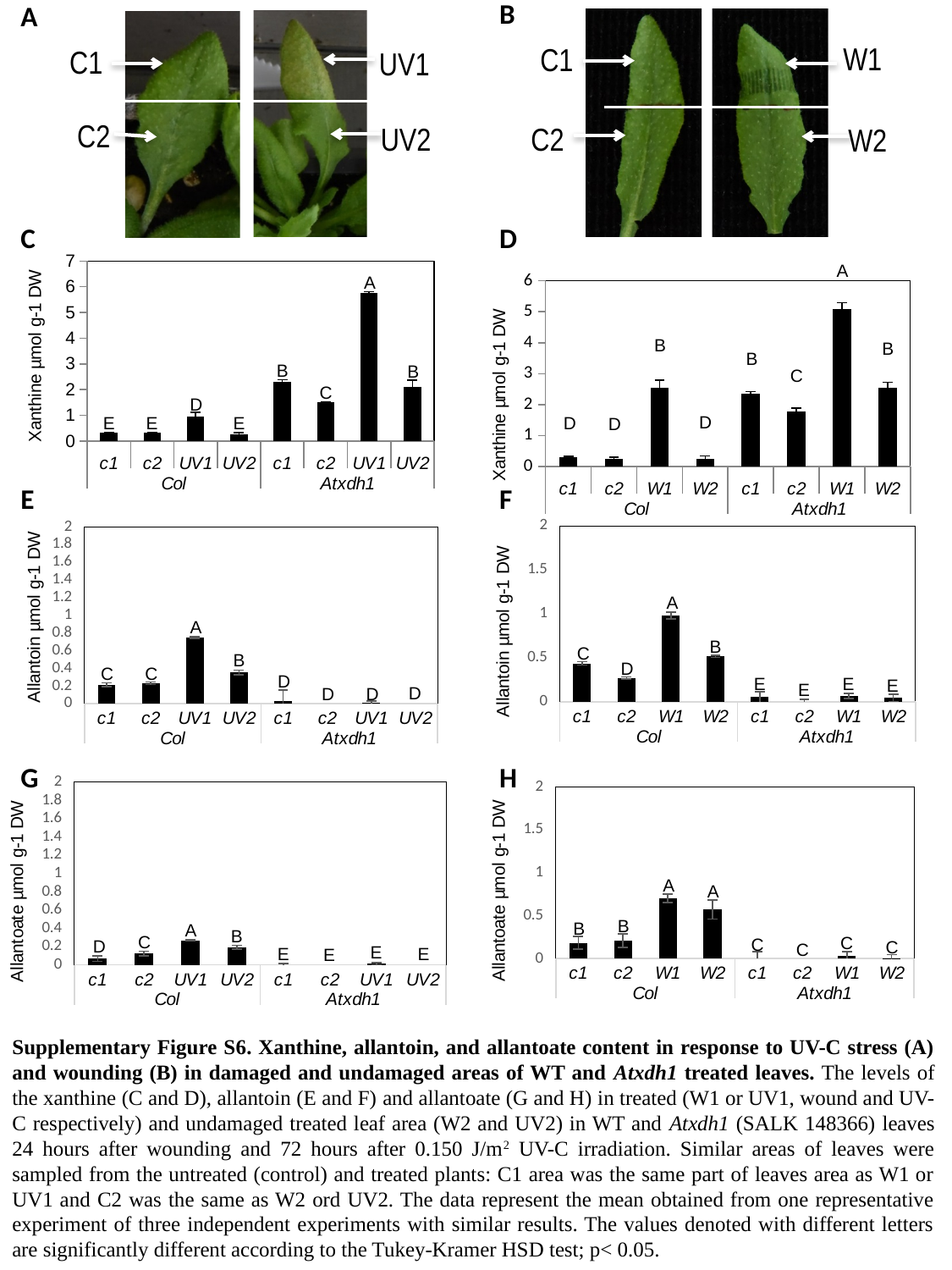

B
A
W1
C1
C1
UV1
C2
UV2
C2
W2
#### Chart
| Category | |
|---|---|
| c1 | 0.300287224730455 |
| c2 | 0.2472953615427276 |
| W1 | 2.5478916037735546 |
| W2 | 0.2447911047169784 |
| c1 | 2.3658993463611586 |
| c2 | 1.767599800117903 |
| W1 | 5.095783207547111 |
| W2 | 2.5574461972877054 |C
D
#### Chart
| Category | |
|---|---|
| c1 | 0.33038806142740057 |
| c2 | 0.33326327398729033 |
| UV1 | 0.9664634945631428 |
| UV2 | 0.24752631204697048 |
| c1 | 2.301006792143956 |
| c2 | 1.5126936940765223 |
| UV1 | 5.765693538384421 |
| UV2 | 2.0995586261199315 |A
A
B
B
B
B
B
C
C
D
D
E
E
D
E
D
E
F
#### Chart
| Category | control |
|---|---|
| c1 | 0.43645521887263017 |
| c2 | 0.27238224201163475 |
| W1 | 0.9821014773760318 |
| W2 | 0.5165323716849745 |
| c1 | 0.05649875705400331 |
| c2 | -0.005154359477162029 |
| W1 | 0.06767761884262113 |
| W2 | 0.046359789385256756 |
#### Chart
| Category | control |
|---|---|
| c1 | 0.21467062597902542 |
| c2 | 0.23280892665512853 |
| UV1 | 0.7489361687719381 |
| UV2 | 0.3536863104966116 |
| c1 | 0.02899912435649643 |
| c2 | -0.07144511658175146 |
| UV1 | 0.010423316805121985 |
| UV2 | -0.03878701638598214 |A
A
B
C
B
D
C
C
D
E
E
E
E
D
D
D
G
H
#### Chart
| Category | control |
|---|---|
| c1 | 0.06821577584427674 |
| c2 | 0.12646848280927603 |
| UV1 | 0.2673771432433928 |
| UV2 | 0.19344367460807255 |
| c1 | -0.02326716402226825 |
| c2 | -0.08535872006674505 |
| UV1 | 0.012283192002850996 |
| UV2 | -0.051194652906744964 |
#### Chart
| Category | control |
|---|---|
| c1 | 0.1841860998429049 |
| c2 | 0.20972302149153632 |
| W1 | 0.7039387163366563 |
| W2 | 0.5723643733335423 |
| c1 | -0.002435529505759426 |
| c2 | -0.09575066638740082 |
| W1 | 0.02808812892216017 |
| W2 | 0.004436515458392742 |A
A
B
B
A
B
C
C
C
D
C
C
E
E
E
E
Supplementary Figure S6. Xanthine, allantoin, and allantoate content in response to UV-C stress (A) and wounding (B) in damaged and undamaged areas of WT and Atxdh1 treated leaves. The levels of the xanthine (C and D), allantoin (E and F) and allantoate (G and H) in treated (W1 or UV1, wound and UV-C respectively) and undamaged treated leaf area (W2 and UV2) in WT and Atxdh1 (SALK 148366) leaves 24 hours after wounding and 72 hours after 0.150 J/m^2^ UV-C irradiation. Similar areas of leaves were sampled from the untreated (control) and treated plants: C1 area was the same part of leaves area as W1 or UV1 and C2 was the same as W2 ord UV2. The data represent the mean obtained from one representative experiment of three independent experiments with similar results. The values denoted with different letters are significantly different according to the Tukey-Kramer HSD test; p< 0.05.

### Slide 7
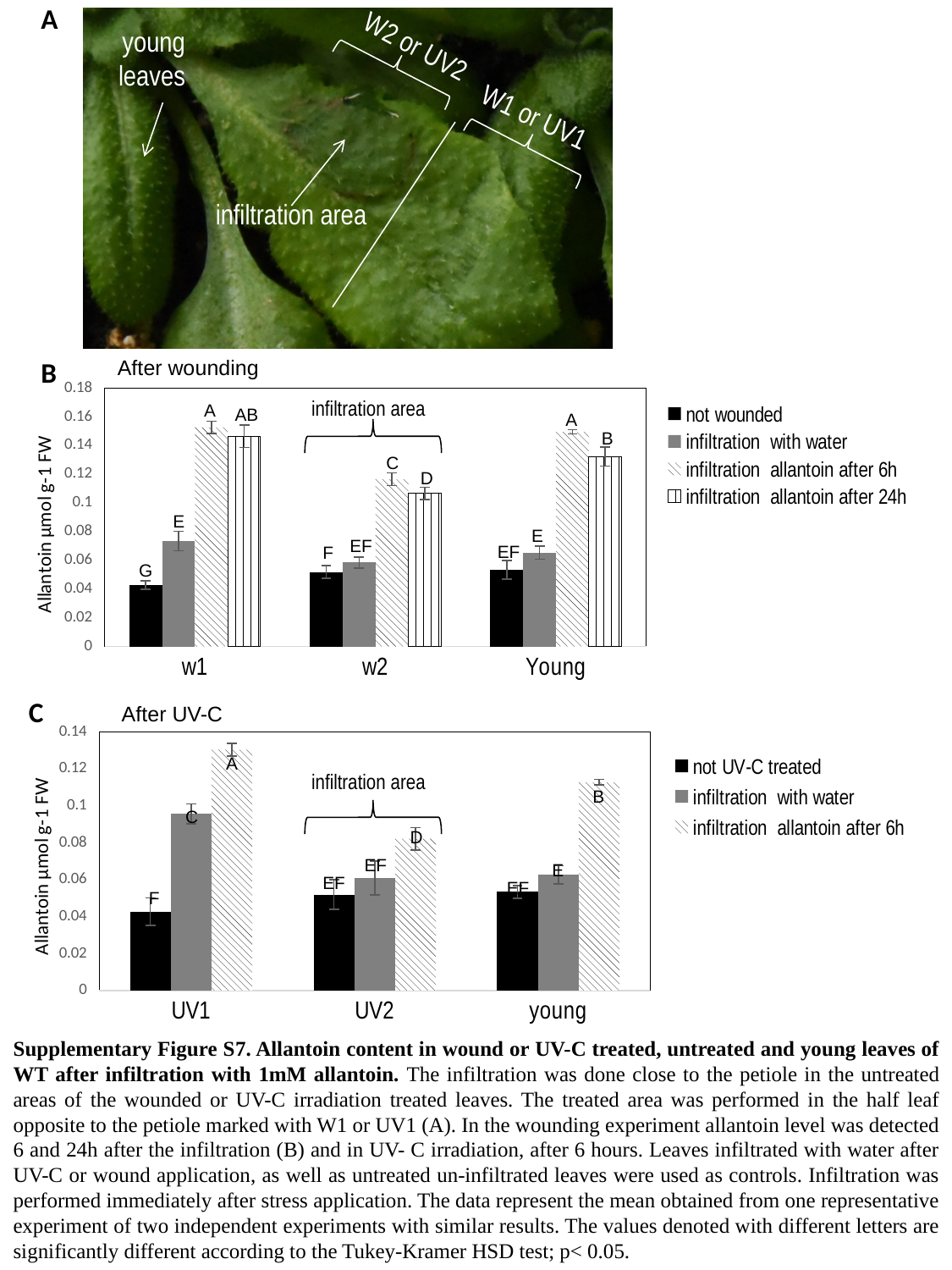

A
W2 or UV2
young leaves
W1 or UV1
infiltration area
After wounding
B
#### Chart
| Category | not wounded | infiltration with water | infiltration allantoin after 6h | infiltration allantoin after 24h |
|---|---|---|---|---|
| w1 | 0.04280607124937281 | 0.07341381108389165 | 0.15264700963175032 | 0.14656235094435466 |
| w2 | 0.051926841986167274 | 0.05853821709315939 | 0.11655267119127473 | 0.10662318113396892 |
| Young | 0.0534164018509227 | 0.06544572671015221 | 0.149446733980918 | 0.13234727342801256 |infiltration area
A
AB
A
B
C
D
E
E
EF
EF
F
G
C
After UV-C
#### Chart
| Category | not UV-C treated | infiltration with water | infiltration allantoin after 6h |
|---|---|---|---|
| UV1 | 0.04280607124937281 | 0.09566817193510621 | 0.13045659809332666 |
| UV2 | 0.051926841986167274 | 0.0609710653955511 | 0.0822271891012483 |
| young | 0.0534164018509227 | 0.0627077778761613 | 0.1128553110489889 |A
infiltration area
B
C
D
EF
E
EF
EF
F
Supplementary Figure S7. Allantoin content in wound or UV-C treated, untreated and young leaves of WT after infiltration with 1mM allantoin. The infiltration was done close to the petiole in the untreated areas of the wounded or UV-C irradiation treated leaves. The treated area was performed in the half leaf opposite to the petiole marked with W1 or UV1 (A). In the wounding experiment allantoin level was detected 6 and 24h after the infiltration (B) and in UV- C irradiation, after 6 hours. Leaves infiltrated with water after UV-C or wound application, as well as untreated un-infiltrated leaves were used as controls. Infiltration was performed immediately after stress application. The data represent the mean obtained from one representative experiment of two independent experiments with similar results. The values denoted with different letters are significantly different according to the Tukey-Kramer HSD test; p< 0.05.

### Slide 8
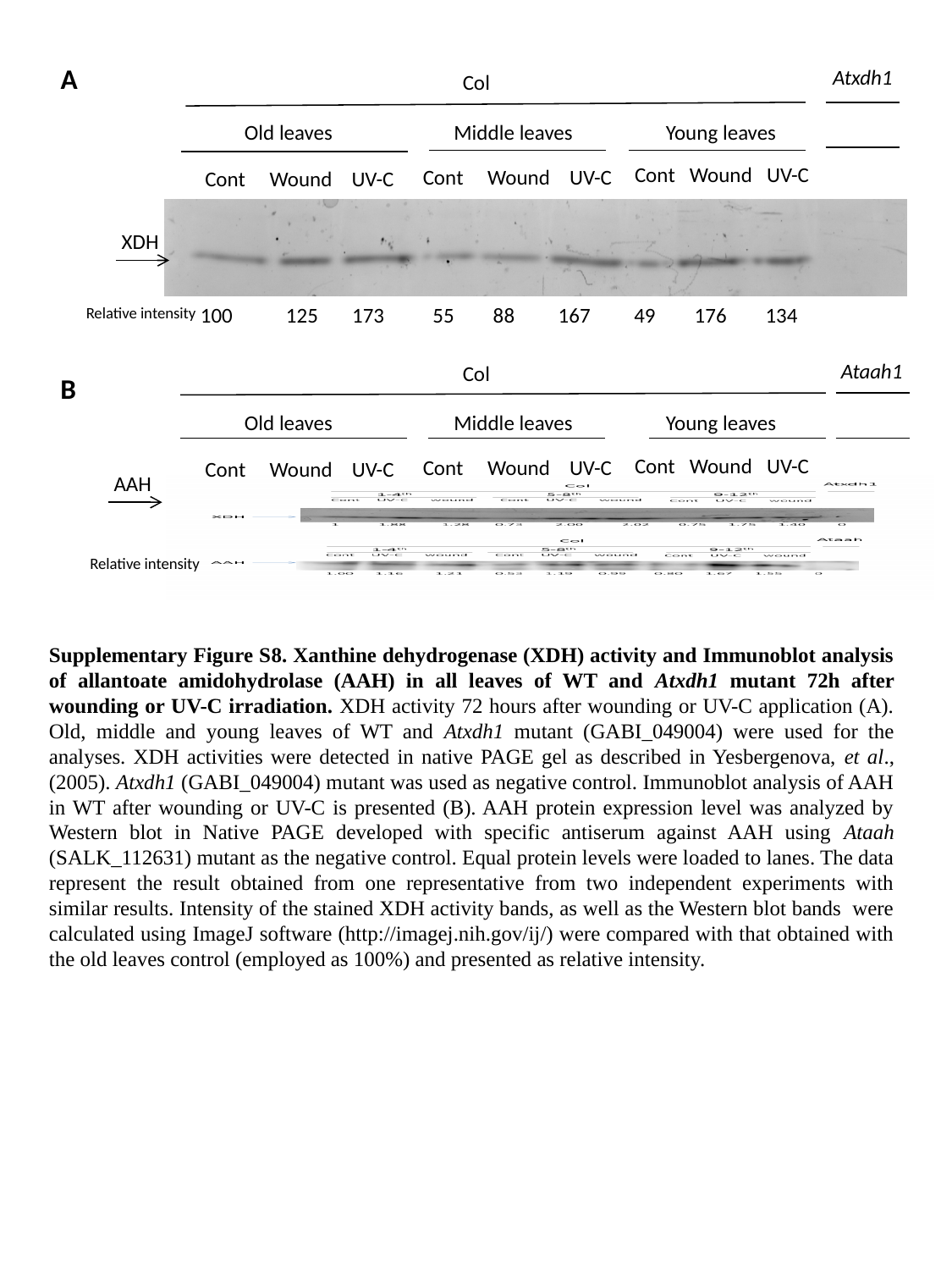

A
Atxdh1
Col
Old leaves
Middle leaves
Young leaves
Cont Wound UV-C
Cont Wound UV-C
Cont Wound UV-C
XDH
100 125 173 55 88 167 49 176 134
Relative intensity
Ataah1
Col
B
Old leaves
Middle leaves
Young leaves
Cont Wound UV-C
Cont Wound UV-C
Cont Wound UV-C
AAH
Relative intensity
Supplementary Figure S8. Xanthine dehydrogenase (XDH) activity and Immunoblot analysis of allantoate amidohydrolase (AAH) in all leaves of WT and Atxdh1 mutant 72h after wounding or UV-C irradiation. XDH activity 72 hours after wounding or UV-C application (A). Old, middle and young leaves of WT and Atxdh1 mutant (GABI_049004) were used for the analyses. XDH activities were detected in native PAGE gel as described in Yesbergenova, et al., (2005). Atxdh1 (GABI_049004) mutant was used as negative control. Immunoblot analysis of AAH in WT after wounding or UV-C is presented (B). AAH protein expression level was analyzed by Western blot in Native PAGE developed with specific antiserum against AAH using Ataah (SALK_112631) mutant as the negative control. Equal protein levels were loaded to lanes. The data represent the result obtained from one representative from two independent experiments with similar results. Intensity of the stained XDH activity bands, as well as the Western blot bands were calculated using ImageJ software (http://imagej.nih.gov/ij/) were compared with that obtained with the old leaves control (employed as 100%) and presented as relative intensity.

### Slide 9
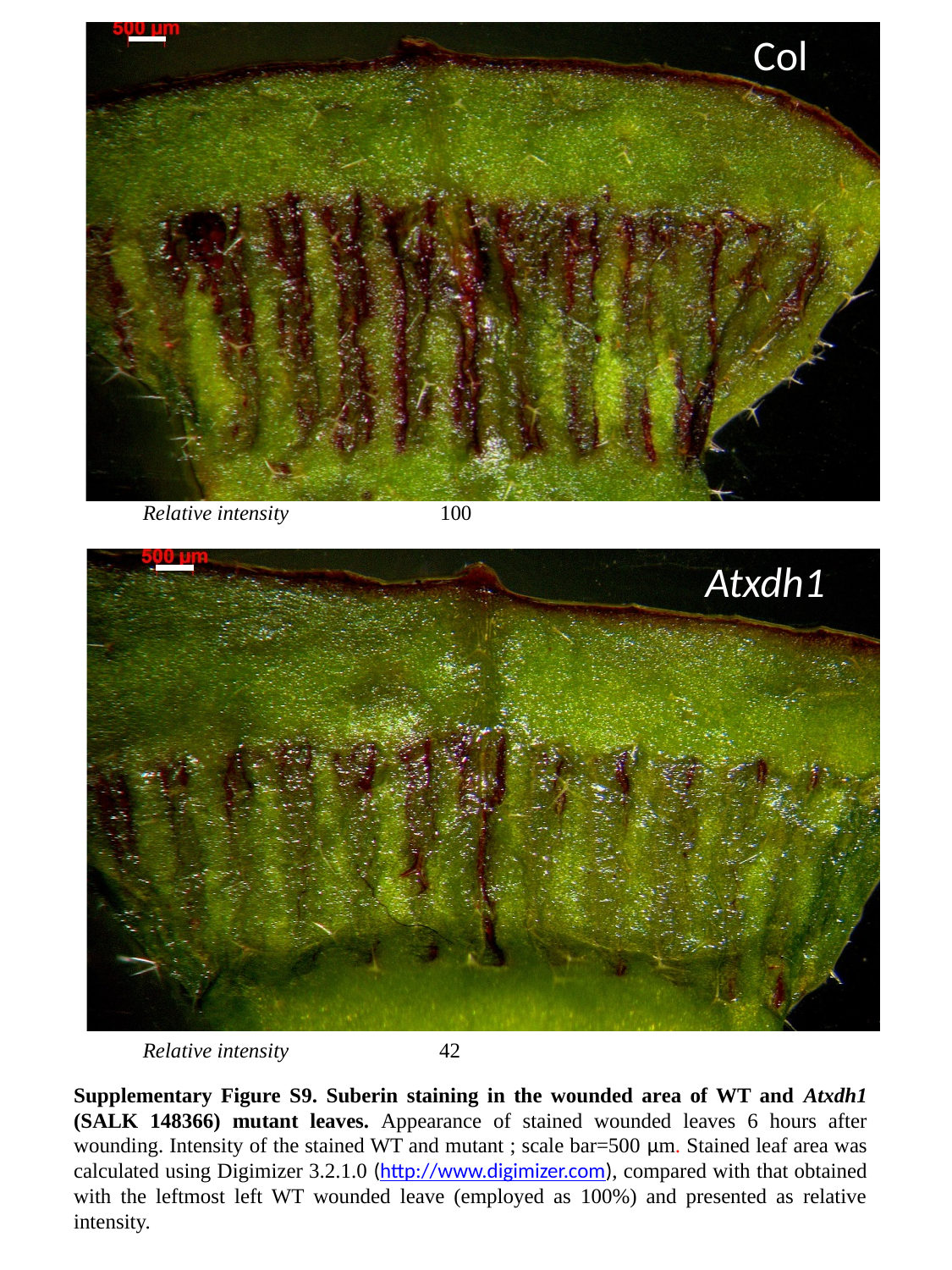

Col
Relative intensity
100
Atxdh1
Relative intensity
42
Supplementary Figure S9. Suberin staining in the wounded area of WT and Atxdh1 (SALK 148366) mutant leaves. Appearance of stained wounded leaves 6 hours after wounding. Intensity of the stained WT and mutant ; scale bar=500 µm. Stained leaf area was calculated using Digimizer 3.2.1.0 (http://www.digimizer.com), compared with that obtained with the leftmost left WT wounded leave (employed as 100%) and presented as relative intensity.

### Slide 10
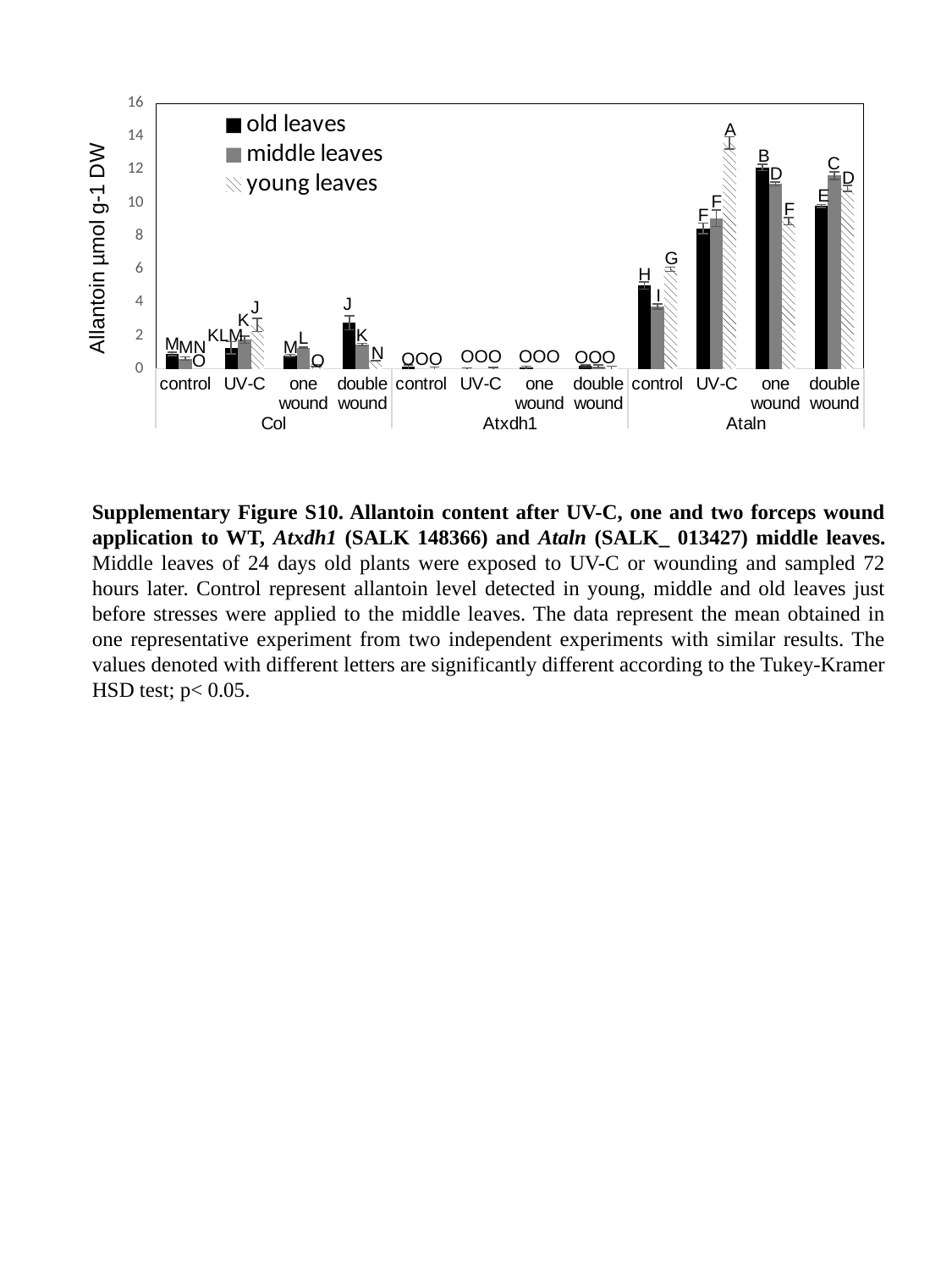

#### Chart
| Category | old leaves | middle leaves | young leaves |
|---|---|---|---|
| control | 0.8998323557672872 | 0.6126122454339693 | -0.12748753947713687 |
| UV-C | 1.260561940127112 | 1.7464505747571655 | 2.6388990873429776 |
| one wound | 0.813333655947378 | 1.2754717822929342 | 0.20249694145074115 |
| double wound | 2.7687252623960434 | 1.4615157004695003 | 0.49319526766516053 |
| control | 0.11294396490361058 | -0.3507011253532607 | -0.018599301372402954 |
| UV-C | -0.038560985171188 | -0.06525774086728331 | -0.013612267830421021 |
| one wound | 0.07115239580506294 | -0.25333250804289387 | -0.253581202479754 |
| double wound | 0.1696996628996279 | 0.1567681167772793 | 0.051091935750973594 |
| control | 5.01428071933205 | 3.7531743599747163 | 6.001146279288731 |
| UV-C | 8.460957223121115 | 9.08163978795651 | 13.614906611364345 |
| one wound | 12.139759337005076 | 11.137775187815876 | 8.887478280764377 |
| double wound | 9.809605765540676 | 11.645226120028978 | 10.871126229108532 |A
B
C
D
D
E
F
F
F
G
H
I
J
J
K
KLM
K
L
M
M
N
OOO
OOO
OOO
OOO
O
O
MN
Supplementary Figure S10. Allantoin content after UV-C, one and two forceps wound application to WT, Atxdh1 (SALK 148366) and Ataln (SALK_ 013427) middle leaves. Middle leaves of 24 days old plants were exposed to UV-C or wounding and sampled 72 hours later. Control represent allantoin level detected in young, middle and old leaves just before stresses were applied to the middle leaves. The data represent the mean obtained in one representative experiment from two independent experiments with similar results. The values denoted with different letters are significantly different according to the Tukey-Kramer HSD test; p< 0.05.

### Slide 11
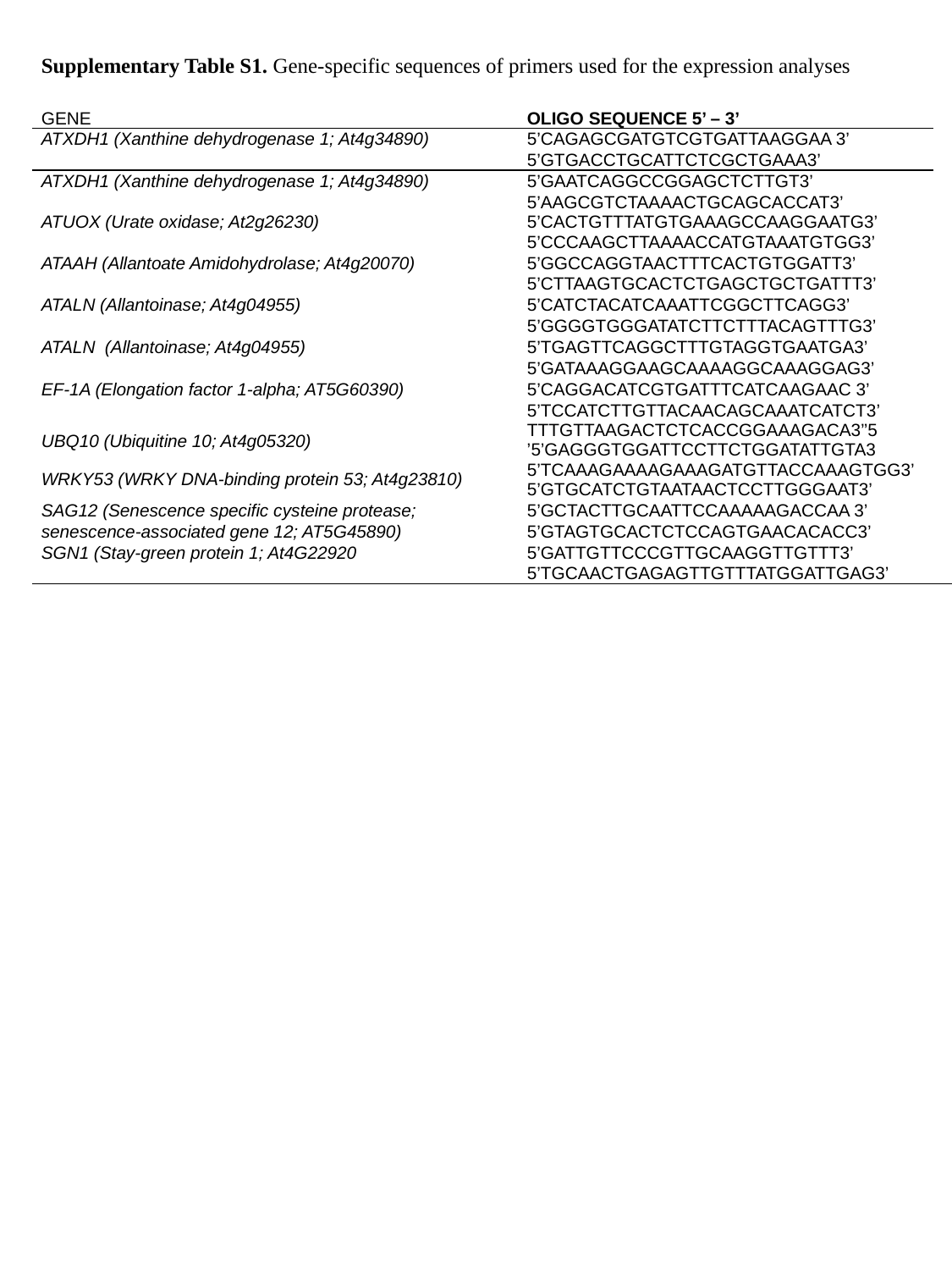

| Supplementary Table S1. Gene-specific sequences of primers used for the expression analyses | |
| --- | --- |
| GENE | OLIGO SEQUENCE 5’ – 3’ |
| ATXDH1 (Xanthine dehydrogenase 1; At4g34890) | 5’CAGAGCGATGTCGTGATTAAGGAA 3’ 5’GTGACCTGCATTCTCGCTGAAA3’ |
| ATXDH1 (Xanthine dehydrogenase 1; At4g34890) | 5’GAATCAGGCCGGAGCTCTTGT3’ 5’AAGCGTCTAAAACTGCAGCACCAT3’ |
| ATUOX (Urate oxidase; At2g26230) | 5’CACTGTTTATGTGAAAGCCAAGGAATG3’ 5’CCCAAGCTTAAAACCATGTAAATGTGG3’ |
| ATAAH (Allantoate Amidohydrolase; At4g20070) | 5’GGCCAGGTAACTTTCACTGTGGATT3’ 5’CTTAAGTGCACTCTGAGCTGCTGATTT3’ |
| ATALN (Allantoinase; At4g04955) | 5’CATCTACATCAAATTCGGCTTCAGG3’ 5’GGGGTGGGATATCTTCTTTACAGTTTG3’ |
| ATALN (Allantoinase; At4g04955) | 5’TGAGTTCAGGCTTTGTAGGTGAATGA3’ 5’GATAAAGGAAGCAAAAGGCAAAGGAG3’ |
| EF-1Α (Elongation factor 1-alpha; AT5G60390) | 5’CAGGACATCGTGATTTCATCAAGAAC 3’ 5’TCCATCTTGTTACAACAGCAAATCATCT3’ |
| UBQ10 (Ubiquitine 10; At4g05320) | 5’TTTGTTAAGACTCTCACCGGAAAGACA3’ 5’GAGGGTGGATTCCTTCTGGATATTGTA3’ |
| WRKY53 (WRKY DNA-binding protein 53; At4g23810) | 5’TCAAAGAAAAGAAAGATGTTACCAAAGTGG3’ 5’GTGCATCTGTAATAACTCCTTGGGAAT3’ |
| SAG12 (Senescence specific cysteine protease; senescence-associated gene 12; AT5G45890) | 5’GCTACTTGCAATTCCAAAAAGACCAA 3’ 5’GTAGTGCACTCTCCAGTGAACACACC3’ |
| SGN1 (Stay-green protein 1; At4G22920 | 5’GATTGTTCCCGTTGCAAGGTTGTTT3’ 5’TGCAACTGAGAGTTGTTTATGGATTGAG3’ |
